# Integrative genomic analysis in African American children with asthma finds 3 novel loci associated with lung function

**DOI:** 10.1101/2020.05.01.045468

**Authors:** Pagé C. Goddard, Kevin L. Keys, Angel C.Y. Mak, Eunice Yujung Lee, Amy K. Liu, Lesly-Anne Samedy-Bates, Oona Risse-Adams, María G. Contreras, Jennifer R. Elhawary, Donglei Hu, Scott Huntsman, Sam S. Oh, Sandra Salazar, Celeste Eng, Blanca E. Himes, Marquitta J. White, Esteban G. Burchard

**Author notes:** Corresponding Author: Kevin L. Keys, PhD, Lung Biology Center, Department of Medicine, UCSF Box 2911, San Francisco, CA 94158 USA. These authors share principal authorship equally (Pagé C. Goddard, Kevin L. Keys). These authors share senior authorship equally (Marquitta J. White, Esteban G. Burchard).

## Abstract

Bronchodilator drugs are commonly prescribed for treatment and management of obstructive lung function present with diseases such as asthma. Administration of bronchodilator medication can partially or fully restore lung function as measured by pulmonary function tests. The genetics of baseline lung function measures taken prior to bronchodilator medication has been extensively studied, and the genetics of the bronchodilator response itself has received some attention. However, few studies have focused on the genetics of post-bronchodilator lung function. To address this gap, we analyzed lung function phenotypes in 1,103 subjects from the Study of African Americans, Asthma, Genes, and Environment (SAGE), a pediatric asthma case-control cohort, using an integrative genomic analysis approach that combined genotype, locus-specific genetic ancestry, and functional annotation information. We integrated genome-wide association study (GWAS) results with an admixture mapping scan of three pulmonary function tests (FEV_1_, FVC, and FEV_1_/FVC) taken before and after albuterol bronchodilator administration on the same subjects, yielding six traits. We identified 18 GWAS loci, and 5 additional loci from admixture mapping, spanning several known and novel lung function candidate genes. Most loci identified via admixture mapping exhibited wide variation in minor allele frequency across genotyped global populations. Functional fine-mapping revealed an enrichment of epigenetic annotations from peripheral blood mononuclear cells, fetal lung tissue, and lung fibroblasts. Our results point to three novel potential genetic drivers of pre- and post-bronchodilator lung function: *ADAMTS1, RAD54B*, and *EGLN3*.

## Introduction

Asthma is a disease characterized by episodic obstruction of airways that affects nearly 339 million people worldwide (The Global Asthma Network, 2018) and is the most common chronic disease among children. Asthma constitutes a massive global economic burden, representing $81.9 billion in medical costs in the United States alone (Nurmagambetov et al., 2018). As a complex disease, asthma results from both environmental and genetic factors, with genetic heritability estimates ranging from 0.35 to 0.90 (Ober & Yao, 2011). The advent of genome-wide association studies (GWAS) (Risch & Merikangas, 1996), combined with progressively larger sample sizes in recent years, has enabled researchers to query the genetic basis of asthma at unprecedented scale, with numerous loci identified in autoimmune and inflammatory pathways (Demenais et al., 2018). However, these loci account for a small portion of asthma liability (Demenais et al., 2018).

Pulmonary function tests are recommended to guide in the diagnosis of asthma and monitor patient status (Asthma and Allergy Foundation of America, 2019). During these tests, patients breathe through a spirometer that captures key measures of lung function, including the forced expiratory volume in 1 second (FEV_1_), which measures initial forced exhalatory capacity; the forced vital capacity (FVC), which measures the maximum total volume of air that a patient can forcibly exhale; and their ratio (FEV_1_/FVC). Lung function measures can be population-normalized according to expected lung function values that account for age, sex, height and ethnicity of the patient (Hankinson et al., 1999). Spirometric measurements can be taken both before bronchodilator treatment (Pre-BD) and after (Post-BD) to further understand lung function status. Historically, baseline lung function is measured with Pre-BD measures, but among people with asthma, post-BD lung function may best reflect lung health (Brehm et al., 2015).

While the genetic contribution to asthma and lung function has been extensively studied via GWAS, most analyses have relied on subjects of European descent (Demenais et al., 2018; Johansson et al., 2019; Pickrell et al., 2016; Z. Zhu et al., 2018). This overrepresentation of ethnically white subjects in biomedical research has impaired the generalizability of genetic studies of complex disease (Burchard, 2014; Bustamante et al., 2011; Popejoy & Fullerton, 2016). Ethnic differences in lung function, particularly between African Americans and European Americans, have been reported for over 40 years (Binder et al., 1976; Glindmeyer et al., 1995; Hsi et al., 1983; Rossiter & Weill, 1974; Schwartz et al., 1988). Ethnic disparities in lung function were attributed to population differences in sitting height, as increased height leads to increased lung capacity. However, adjustment for sitting height only explains 42-50% of ethnic differences in lung function between African Americans and European Americans (Harik-Khan et al., 2004), suggesting that a simplistic reduction to ethnic differences in height cannot account for the observed disparity in lung function. Unequal socioeconomic conditions were also thought to contribute to ethnic differences in lung function (Braun, 2015; Quanjer, 2013, 2015), but socioeconomic factors only account for 7-10% of unexplained variance (Harik-Khan et al., 2004). Self-identified race or ethnicity are commonly used in the clinic to interpret lung function measures, but these are not ideal variables for understanding *genetic* differences in lung function between populations. Kumar *et al.* observed that the proportion of global African genetic ancestry is inversely correlated with lung function (Kumar et al., 2010). Spear *et al.* later observed population differences among African Americans, Mexican Americans, and Puerto Ricans in bronchodilator drug response to albuterol, the short-acting β_2_-adrenergic receptor agonist that is the most commonly prescribed drug for the treatment of acute asthma symptoms (Spear et al., 2019). Specifically, Spear *et al*. performed admixture mapping, a technique designed to identify regions of the genome where locus-specific ancestry drives variation in a disease trait (Shriner, 2013) that has been helpful in studies of complex diseases, including asthma and breast cancer (Féjerman et al., 2012; Pino-Yanes et al., 2015). However, admixture mapping studies comparing baseline and post-bronchodilator lung function have not yet been performed in African Americans. In this study, we address this gap in knowledge by evaluating the effect of locus-specific ancestry on both pre- and post-bronchodilator lung function measures in a pediatric case-control cohort of African Americans children and adolescents.

## Methods

### Study Population

The Study of African Americans, Asthma, Genes and Environments (SAGE) is a case-control cross-sectional cohort study of genetics and gene-environment interactions in African American children and adolescents in the USA. SAGE includes detailed clinical, social, and environmental data on both asthma and asthma-related conditions. Full details of the SAGE study protocols are described in detail elsewhere (Borrell et al., 2013; Nishimura et al., 2013; Thakur et al., 2013; Mak et al., 2018). Briefly, SAGE was initiated in 2006 and recruited participants with and without asthma through a combination of clinic- and community-based recruitment centers in the San Francisco Bay Area. All participants in SAGE self-identified as African American and self-reported that all four grandparents were African American.

Pulmonary function tests were taken prior to administration of albuterol bronchodilator medication for all individuals, both those with and without asthma. Post-bronchodilator spirometry measures were performed only for individuals with asthma. Analyses of pre-bronchodilator lung function measures included all 1,103 asthma cases and controls with complete covariate information. Post-bronchodilator analyses were performed on the 831 asthma cases with post-bronchodilator measurements.

### Genotyping and Quality Control

DNA was isolated from whole blood collected from SAGE participants at the time of study enrollment as described previously (Borrell et al., 2013). DNA was extracted using the Wizard® Genomic DNA Purification kits (Promega, Fitchburg, WI). Samples were genotyped with the Affymetrix Axiom LAT1 array (World Array 4, Affymetrix, Santa Clara, CA).

Genotype quality control was performed in PLINK v1.9 (Chang et al., 2015). Of the 772,703 genotyped variants, 111,901 SNPs were excluded from analysis due to genotype missingness more than 5% (n = 28,211), minor allele frequency (MAF) less than 1% (n = 80,420) or deviation from Hardy Weinberg expectations (HWE) at *p* < 0.001 (n = 3,270). The final set of 660,802 genotyped markers. (Supplementary Table 1).

Genotyped SNPs were submitted to the Michigan Imputation Server (Das et al., 2016), phased using EAGLE v2.3 (Loh et al., 2016), and imputed from the 1000 Genomes Project reference panel (The 1000 Genomes Project Consortium, 2015) using Minimac3 (Das et al., 2016). Imputed SNPs with imputation *R*^2^ < 0.3, with deviation from Hardy-Weinberg equilibrium (HWE) *p*-value < 10^−4^, or with minor allele frequency (MAF) < 1% were discarded. Of the 47,101,126 imputed SNPs, a total of 31,146,322 were culled due to either low MAF (n = 31,095,418) or deviation from HWE (n = 50,904). All variants in the imputed set showed a genotype missingness of no more than 5%. The final number of SNPs used in association analyses was 15,954,804 (Supplementary Table 1).

### Outcome Phenotypes

Pulmonary function testing was performed at the time of recruitment according to the American Thoracic Society / European Respiratory Society standards (Miller et al., 2005; Pellegrino et al., 2005; Wanger et al., 2005) with a KoKo PFT Spirometer (nSpire Health Inc., Louisville, CO). Spirometry was performed both before and 15 minutes after administration of four puffs of albuterol (90ug per puff) through a 5-cm plastic mouthpiece from a standard metered-dose inhaler. Patients were assessed for the following spirometric measures before and after bronchodilator drug usage (pre-BD and post-BD, respectively): (a) FEV_1_, (b) FVC, and (c) FEV_1_/FVC. A total of six phenotypes were assessed for genotype association: pre-BD FEV_1_ (Pre-FEV_1_), pre-BD FVC (Pre-FVC), pre-BD FEV_1_/FVC (Pre-FEV_1_/FVC), post-BD FEV_1_ (Post-FEV_1_), post-BD FVC (Post-FVC), and post-BD FEV_1_/FVC (Post-FEV_1_/FVC). All phenotype values were normalized based on the expected lung function values calculated from the Hankinson equations (Hankinson et al., 1999), which account for age, sex, height, and self-reported ethnicity. Phenotype distributions were checked for normality and to detect outliers. Outliers were determined using the method of Tukey fences (John Tukey, 1977). For each phenotype, we computed the first quartile value (Q1), the third quartile value (Q3), and the interquartile range (IQR). We declared as outliers all values outside of the range 

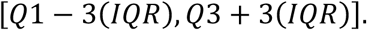

Individuals with outlier values for a phenotype were removed from association analyses for that phenotype.

### Covariates

#### Age, Sex, and BMI

Biometric covariates such as age, sex, body mass index (BMI), and height were measured directly at time of recruitment. BMI was categorized into underweight, normal, overweight, and obese, according to CDC guidelines for defining childhood obesity (Barlow, 2007; Cote et al., 2013; Whitlock et al., 2005). An overweight status was defined as a BMI at or above the 85th percentile for the general population of children of the same sex and in the same age group. An obese status was defined as a BMI at or above the 95th percentile. Underweight individuals (bottom 5th percentile, n = 9) were excluded from analysis.

#### Asthma status

Case status was defined as physician-diagnosed asthma supported by reported asthma medication use and symptoms of coughing, wheezing, or shortness of breath in the 2 years preceding enrollment.

#### Maternal Educational Attainment

Maternal educational attainment was measured at recruitment and included in analyses to control for socioeconomic status. It was coded as total years of education completed from the first grade: for example, a complete K-6 education was 6 years, a complete high school education was 12 years, and any additional years (college or trade school and beyond) were counted as 1 year each.

#### Genetic Ancestry

Global genetic ancestry was estimated for each individual with the ADMIXTURE software (Alexander et al., 2009) in supervised learning mode assuming one West African and one European ancestral population, with HapMap Phase III YRI and CEU populations as references (The International HapMap 3 Consortium, 2010). Local ancestry estimation was performed with RFMix (Maples et al., 2013; Spear et al., 2019) using the same two-way ancestry reference from HapMap Phase III.

#### Estimation of Genetic Relatedness and Genotype Principal Components

Genetic relatedness matrices (GRMs) were generated in R using GENESIS (Gogarten et al., 2019), which provides a computational pipeline for handling complex population structure. We used PCAir (Conomos et al., 2015) to correct for distant population structure accounting for relatedness, and PC-Relate (Conomos et al., 2016) to adjust for genetic relatedness in recently admixed populations. The resulting principal components provide better correction for population stratification in admixed populations compared to standard PCA on genotypes (Patterson et al., 2006).

### Genetic Association Analyses

Genotype association testing was performed with the MLMA-LOCO algorithm from GCTA (Yang et al., 2011, 2014) to correct for population structure using GRMs generated with GENESIS. Association of outcome phenotypes with allele dosages at 15,954,804 biallelic SNPs was performed with a “leave one chromosome out” model to avoid double-fitting tested variants. Other variables included in models were age, sex, BMI, maternal educational attainment, and three genotype principal components. Models of Pre-BD also included asthma status.

The suggestive and significant association thresholds for each outcome phenotype were determined by the effective number of independent statistical tests (M_eff_) calculated with CODA (Plummer et al., 2006). CODA computes M_eff_ using the autocorrelation of *p*-values from GWAS. For our analyses, M_eff_ ranged from 488,819 to 507,975 (Supplementary Table 2). The Bonferroni corrected genome-wide significant threshold was computed as 0.05/M_eff_, while the suggestive threshold was computed as (1/M_eff_), yielding a single pair of thresholds for all six outcome phenotypes considered: *p* < 1.99 × 10^−6^ for suggestive association, and *p* < 9.95 × 10^−8^ for significant association (Supplementary Table 2).

Admixture mapping analyses were performed using linear regression models in R and local ancestry calls from RFMix for 454,322 genotyped SNPs. Counts of 0, 1, or 2 alleles of African descent were computed for each person at each SNP. Phenotypes were then regressed onto ancestral allele counts for each SNP while including age, sex, height, BMI, maternal educational attainment (as a proxy for socioeconomic status), and global African genetic ancestry proportion as covariates. Analyses with pre-BD outcome measures also included asthma status as a covariate.

### Fine-mapping genetic associations

Functional fine-mapping with PAINTOR (Kichaev et al., 2014) was used to identify putative causal variants in novel loci deemed statistically significant by admixture mapping. PAINTOR applies a Bayesian probabilistic framework to integrate functional annotations, association summary statistics (Z-scores), and linkage disequilibrium information for each locus to prioritize the most likely causal variants in a given region. Functional annotations were selected per locus as recommended by the authors of PAINTOR (Kichaev, 2017). A subset of lung- and blood-related functional annotations from the Roadmap Epigenomics Project (Roadmap Epigenomics Consortium et al., 2015) and the ENCODE Consortium (ENCODE Project Consortium, 2012) were assessed for their individual improvement to the posterior probability of causality; the top 5 minimally correlated annotations were selected for each locus.

### Annotation Tools

The NHGRI/EBI GWAS Catalog (Buniello et al., 2019), Ensembl Genome Browser release 98 (Cunningham et al., 2019) and gnomAD browser v3.0 (Karczewski et al., 2019) were used to look up known associations at significant loci according to our analyses. Annotation lookups in the gnomAD browser v3.0 used hg38 coordinates translated from our hg19-aligned genotypes via liftOver (Hinrichs et al., 2006). Data management, statistical analysis, and figure generation made extensive use of GNU parallel (Tange, 2018) and several R packages, including data.table, doParallel, optparse, ggplot2, and the tidyverse bundle (Calaway et al., 2018; Davis et al., 2019; Dowle et al., 2019; Wickham, Hadley, 2016, p. 2; Wickham, Hadley & Grolemund, Garrett, 2017).

## Results

### Cohort Characteristics

Characteristics of all SAGE participants included in analyses are shown in Table 1. Distributions of each lung function measure stratified by case/control status and bronchodilator administration (pre-BD vs. post-BD) are shown in Supplementary Figure 1. FVC showed no significant difference between asthma cases and controls (Kruskal-Wallis *p*-value = 0.073), while stratification by case/control status yielded significantly different distributions for FEV_1_ (Kruskal-Wallis *p*-value = 4.8 x 10-7) and FEV_1_/FVC (Kruskal-Wallis *p*-value = 1.5 x 10-7). Among cases, statistically significant differences were observed between distributions of pre-BD and post-BD measures of FEV_1_ (Kruskal-Wallis *p*-value = 1.2 x 10-38), FVC (Kruskal-Wallis *p*-value = 5.4 x 10-16), and FEV_1_/FVC (Kruskal-Wallis *p*-value = 4.0 x 10^−29^), illustrating a measurable effect of bronchodilator medication on lung function.

**Table 1:**
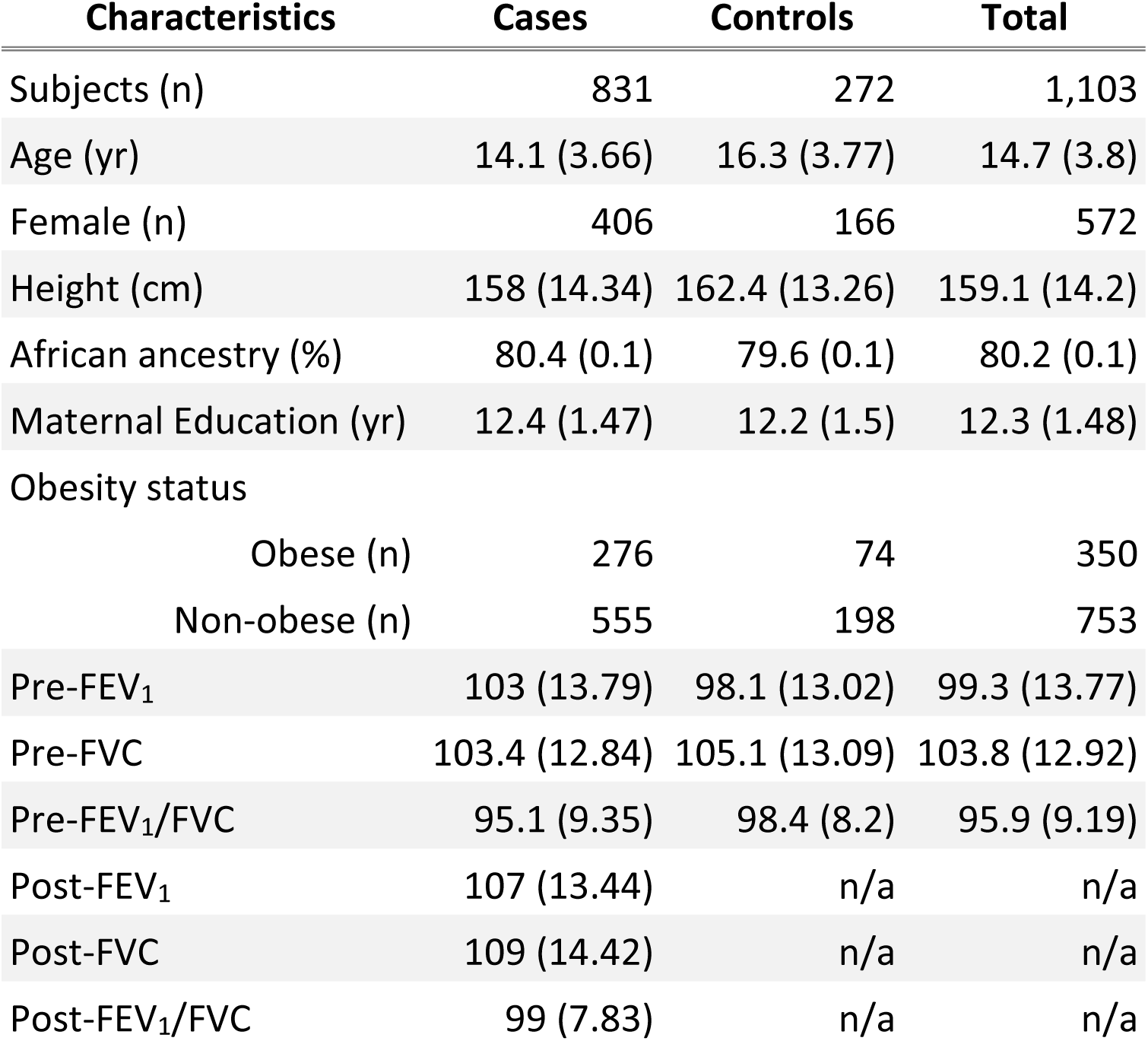
Summary statistics of phenotypes and covariates from the SAGE cohort. Displayed numbers are either counts (n) or averages followed by standard errors in parentheses. Units are listed where appropriate. An “n/a” appears where measurements taken on cases only.

| Characteristics | Cases | Controls | Total |
| --- | --- | --- | --- |
| Subjects (n) | 831 | 272 | 1,103 |
| Age (yr) | 14.1 (3.66) | 16.3 (3.77) | 14.7 (3.8) |
| Female (n) | 406 | 166 | 572 |
| Height (cm) | 158 (14.34) | 162.4 (13.26) | 159.1 (14.2) |
| African ancestry (%) | 80.4 (0.1) | 79.6 (0.1) | 80.2 (0.1) |
| Maternal Education (yr) | 12.4 (1.47) | 12.2 (1.5) | 12.3 (1.48) |
| Obesity status |  |  |  |
| Obese (n) | 276 | 74 | 350 |
| Non-obese (n) | 555 | 198 | 753 |
| Pre-FEV<sub>1</sub> | 103 (13.79) | 98.1 (13.02) | 99.3 (13.77) |
| Pre-FVC | 103.4 (12.84) | 105.1 (13.09) | 103.8 (12.92) |
| Pre-FEV<sub>1</sub>/FVC | 95.1 (9.35) | 98.4 (8.2) | 95.9 (9.19) |
| Post-FEV<sub>1</sub> | 107 (13.44) | n/a | n/a |
| Post-FVC | 109 (14.42) | n/a | n/a |
| Post-FEV<sub>1</sub>/FVC | 99 (7.83) | n/a | n/a |

The global African genetic ancestry proportion in our sample varied from 30.7% to 100%, with an average proportion of 80.2% (Supplementary Figure 2), concordant with empirically observed averages (Baharian et al., 2016). Global ancestry contained the same information as the first genotype principal component (*R*^2^ = 0.984, Supplementary Figure 3).

### Genetic association testing finds novel and known loci

Figure 1 shows results from GWAS performed on pre-BD phenotypes (Pre-FEV_1_, Pre-FVC, and Pre-FEV_1_/FVC) and post-BD phenotypes (Post-FEV_1_, Post-FVC, and Post-FEV_1_/FVC) using linear mixed modeling. The association results showed no evidence of genomic inflation, with genetic control *λ* ranging from 0.98 to 0.99 (Supplementary Table 2, Supplementary Figure 4). Table 2 lists the 18 genome-wide significant associations found, each associated with exactly one of the six lung function measures. An additional 252 variants were suggestively associated with at least one phenotype (Supplementary Tables 3–8). Of the 18 variants, 4 variants on chromosome 13 in a region spanned by the gene *ATP8A2* were associated with Pre-FEV_1_/FVC (Supplementary Figure 6). Two variants on chromosome 16 that were associated with Pre-FVC flanked the promoter region of *IRX3* (Supplementary Figure 7). A third variant associated with Pre-FVC was located on chromosome 20 near *THBD* (Supplementary Figure 8), a gene linked to venous thromboembolism in African American and Afro-Caribbean individuals (Hernandez et al., 2016). Two variants associated with Post-FVC were in a gene-rich region on chromosome 19 (Supplementary Figure 9), with the peak near *TMIGD2* and *SHD*, while eight other variants pointed to a second gene-rich region on chromosome 11 near *CXCR5* and *HYOU1* (Supplementary Figure 10). Post-FEV_1_/FVC was associated with a region on chromosome 15 near the genes *AKAP13* and *ADAMTS7P4* (Supplementary Figure 11).

**Table 2:**
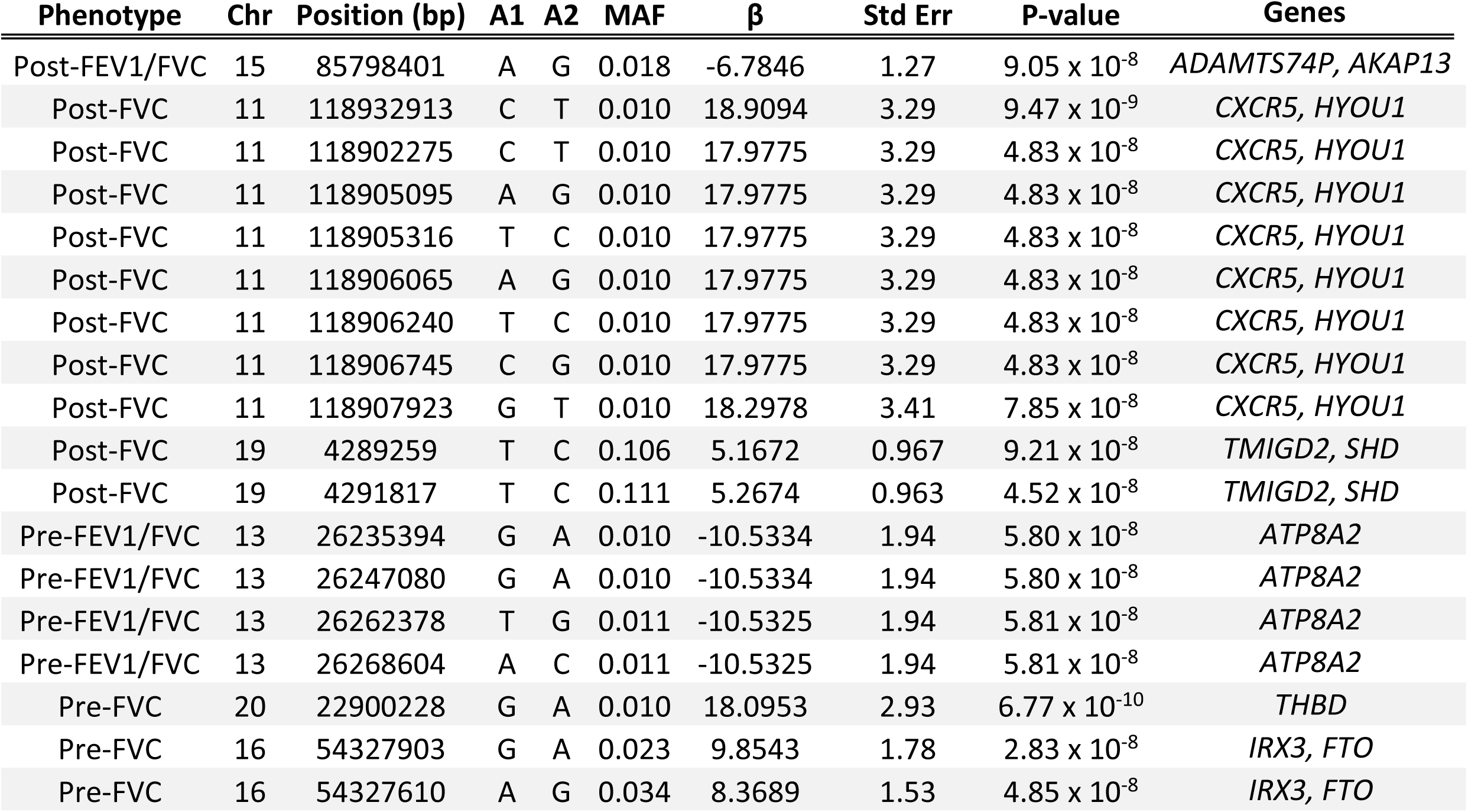
Significant association results from GWAS. The p-values for all SNPs listed here met the significance threshold of 9.95 × 10^−8^. SNPs were specified by chromosome (Chr) and physical position in base pairs (bp). A1 denotes the major allele, while A2 denotes the minor allele. MAF denotes the minor allele frequency (MAF). B denotes the effect size. “Std Err” is the standard error of the estimate of β. “Genes” denotes any genes within proximity of the associated variants.

**Figure 1:**
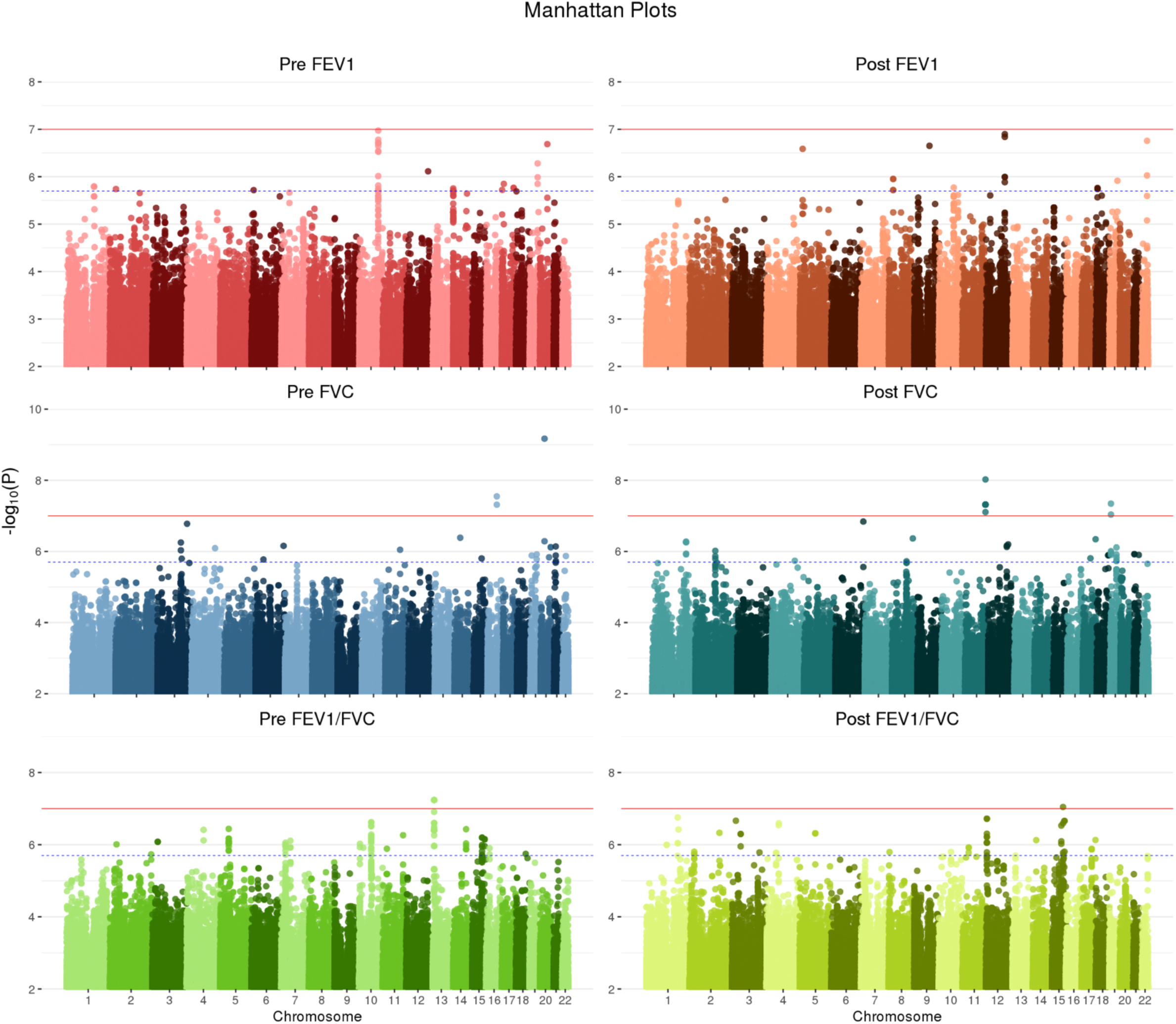
Manhattan plots summarizing GWAS *p*-values for all six lung function phenotypes. The solid red line denotes genome-wide significance (*p*-value < 9.95 × 10^−8^), while the dashed blue line marks the suggestive threshold (*p*-value < 1.99 × 10^−6^), per CODA calculations. Variants with a p-value greater than 0.05 were deemed uninformative and therefore not plotted.

Among the suggestive associations, a variant on chromosome 12 associated with Post-FEV1 was near *BTBD11* (Supplementary Figure 12), a gene previously associated with Post-FEV_1_, Post-FEV_1_/FVC, and ΔFEV_1_, the change in lung function due to bronchodilator administration (Hardin et al., 2016; Lutz et al., 2015), as well as BMI (Kichaev et al., 2019). A suggestive association with Pre-FEV_1_ on chromosome 12 fell near *SCARB1* (Supplementary Figure 13), which was previously associated with FEV_1_ and FVC (Wyss et al., 2018) and HDL cholesterol levels (Wojcik et al., 2019). Another suggestive association with Pre-FEV_1_ on chromosome 20 was near the gene *PTPRT* (Supplementary Figure 14), which was previously associated with thromboembolism susceptibility in 5,334 African American individuals (Heit et al., 2017).

### Admixture mapping identified five novel loci not found by GWAS

Table 3 shows five regions where admixture proportions were statistically significantly associated with one of the six phenotypes. The three pre-BD phenotypes (Pre-FEV_1_, Pre-FVC, Pre-FEV_1_/FVC) were each associated with one region, while Post-FVC was associated with two distinct regions. Post-FEV_1_ and Post-FEV_1_/FVC had no significant associations. None of the regions overlapped with those significant in our GWAS, and none showed large deviations from mean genome-wide African genetic ancestry. A small region on chromosome 21 that was significantly associated with Pre-FEV_1_ flanked the genes *ADAMTS1* and *ADAMTS5* (Supplementary Figure 15). The region on chromosome 4 associated with Pre-FVC pointed to two candidate genes, *RCHY1* and *THAP6*, that had no prior lung disease associations (Supplementary Figure 17). A region on chromosome 19 associated with Pre-FEV_1_/FVC spanned the genes *ZNF557* and *INSR* (Supplementary Figure 18). Post-FVC was associated with two regions, one on chromosome 8 spanning the genes *ESRP1, INTS8, TP53INP1* and *NDUFAF6* (Supplementary Figure 19), and another on chromosome 14 encompassing *EGLN3* and *SNX6* (Supplementary Figure 20).

**Table 3:** Admixture mapping in SAGE identified five regions with statistically significant association to at least one phenotype. The regions are arbitrarily numbered from 1 to 5 and defined by phenotype, chromosome (Chr), physical starting point (in base pairs), and end point (in base pairs). Physical positions are given in hg19 coordinates. The total length of the region is given in kilobasepairs (Kb). The threshold for statistical significance is given for each region. SNP_Admix_ counts the number (n) of genotyped SNPs from admixture mapping that met the significance threshold. SNP_GWAS_ counts the total number of GWAS SNPs (genotyped and/or imputed) in the admixture mapping region. AFR denotes the percentage (%) of local genetic ancestry of African origin. The column “Genes” lists genes physically within and near the associated regions.

| Locus | Phenotype | Chr | Start (bp) | End (bp) | Length (Kb) | Threshold | SNP<sub>Admix</sub> (n) | SNP<sub>GWAS</sub> (n) | AFR (%) | Genes |
| --- | --- | --- | --- | --- | --- | --- | --- | --- | --- | --- |
| 1 | Pre-FEV1 | 21 | 28,209,667 | 28,240,392 | 30.72 | 1.03 x 10<sup>-4</sup> | 4 | 215 | 78.7 | ADAMTS1 |
| 2 | Pre-FVC | 4 | 75,555,658 | 76,873,740 | 1318.08 | 9.64 x 10<sup>-5</sup> | 102 | 7905 | 79.8 | RCHY1, THAP6 |
| 3 | Pre-FEV1/FVC | 19 | 7,068,207 | 7,127,294 | 59.09 | 1.04 x 10<sup>-4</sup> | 18 | 376 | 80.4 | INSR, ZNF557 |
| 4 | Post-FVC | 8 | 95,387,941 | 95,820,594 | 432.65 | 9.93 x 10<sup>-5</sup> | 45 | 2822 | 80.8 | ESRP1, INTS8, TP53INP1, NDUFAF6 |
| 5 | Post-FVC | 14 | 34,283,561 | 34,595,061 | 311.50 | 9.93 x 10<sup>-5</sup> | 95 | 1982 | 80.5 | EGLN3, SNX6 |

### Functional fine-mapping found three novel putatively causal loci for lung function phenotypes

Table 4 lists the most probable causal SNP for each of the five admixture mapping loci according to PAINTOR. SNP rs13615 showed the highest probability of causality (0.630) with Pre-FEV_1_ on locus 1 (Figure 2). This variant falls within the 3’ untranslated region of *ADAMTS1*, suggesting that *ADAMTS1* drives the admixture mapping association and not its physical neighbor *ADAMTS5*. The minor allele frequency of rs13615 in African and African diaspora populations was lower than every other global population (2.6% AFR vs. 7.0 – 54.5% other populations, gnomAD v3; see Supplementary Figure 21). The SNP rs10857225 emerged as the most likely causal variant (probability 0.361) for the association of Pre-FVC with locus 2 on chromosome 4 (Figure 3). This variant is located within an intron of the gene *THAP6*, suggesting that *THAP6* is more likely the causal gene behind the association with Pre-FVC. In contrast to locus 1, the minor allele frequency of rs10857225 is highest in global African populations and markedly lower in other global populations (59.1% AFR vs. 28.1 – 38.4% other populations, gnomAD v3). Locus 3 on chromosome 19 associated with Pre-FEV_1_/FVC, and locus 4 on chromosome 8 associated with Post-FVC, showed little information gain from functional fine-mapping. The driving variant for locus 3, SNP rs72986681, was located in the 3’ untranslated region of *ZNG557*, but showed a low probability of causality (0.168, Supplementary Figure 22). The most probable marker for locus 4, the SNP rs2470740, which is located in intron 2 of *RAD54B*, showed an even lower probability of causality (0.109, Supplementary Figure 23). Functional fine-mapping of locus 5, a region on chromosome 14 associated with Post-FVC, yielded the SNP rs1351618 with a moderate probability of causality (0.390, Figure 4). rs1351618 is located in an intron of *EGLN3*. As with locus 2, rs1351618 had a much higher minor allele frequency in populations of African ancestry versus other global populations (12.4% AFR vs. < 2.2% other populations, gnomAD v3).

**Table 4:** Results from PAINTOR highlighting the most probably causal SNPs for each locus as defined by admixture mapping. Similar to Table 3, the loci are arbitrarily numbered from 1 to 5 and defined by phenotype and chromosome (Chr). The physical position (in basepairs) of the most likely causal SNP is given in hg19 coordinates. Minor allele frequencies (MAF) are taken from global populations from the gnomAD server v3. The displayed *p*-values are from our discovery GWAS. The probability of causality (Pr(causal)) is computed from PAINTOR.

| Locus | Phenotype | Chr | Position (bp) | SNP | Ref allele | Alt allele | MAF (%) | P-value | Annotation | Pr(causal) |
| --- | --- | --- | --- | --- | --- | --- | --- | --- | --- | --- |
| 1 | Pre-FEV<sub>1</sub> | 21 | 28,209,667 | rs13615 | A | G | 2.55% | 6.95 x 10<sup>-3</sup> | ADAMTS1, 3' UTR | 0.630 |
| 2 | Pre-FVC | 4 | 76,464,584 | rs10857225 | A | C | 59.07% | 3.11 x 10<sup>-6</sup> | THAP6, Intron Variant | 0.361 |
| 3 | Pre-FEV<sub>1</sub>/FVC | 19 | 7,087,789 | rs72986681 | G | A | 1.54% | 7.25 x 10<sup>-4</sup> | ZNF557, 3' UTR | 0.168 |
| 4 | Post-FVC | 8 | 95,399,551 | rs2470740 | A | T | 23.82% | 2.04 x 10<sup>-3</sup> | RAD54B, Intron Variant | 0.109 |
| 5 | Post-FVC | 14 | 34,531,633 | rs1351618 | C | T | 12.40% | 1.99 x 10<sup>-4</sup> | EGLN3, Intron Variant | 0.390 |

**Figure 2:**
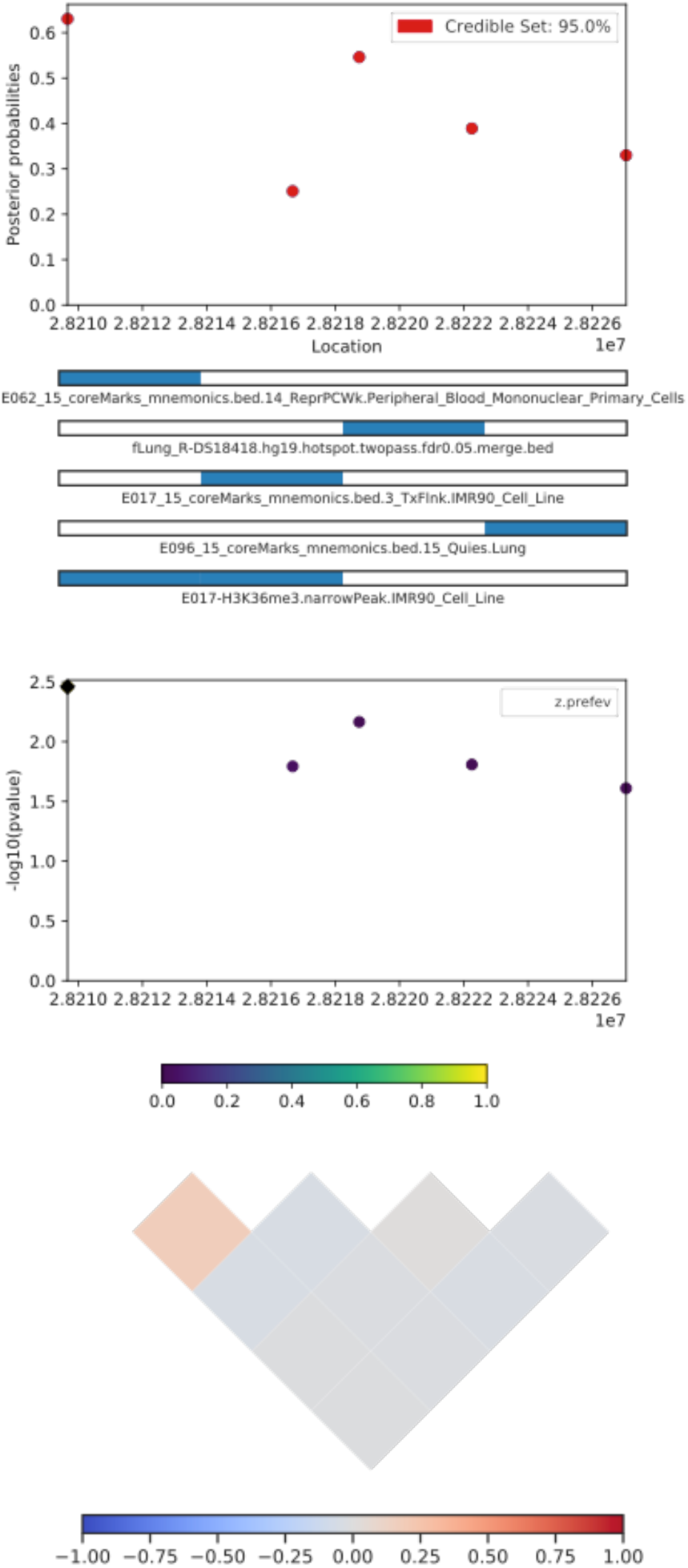
A CANVIS plot of results from PAINTOR functional fine-mapping for locus 1, an association with Pre-FEV_1_ on chromosome 21. The SNP rs13615, which sits in the 3’ UTR of the gene *ADAMTS1*, attains a posterior probability of causality of 0.630. The panels show, from top to bottom, the posterior probability of causality; the 5 most informative functional annotations; GWAS p-values; and local linkage disequilibrium expressed as a signed Pearson correlation.

**Figure 3:**
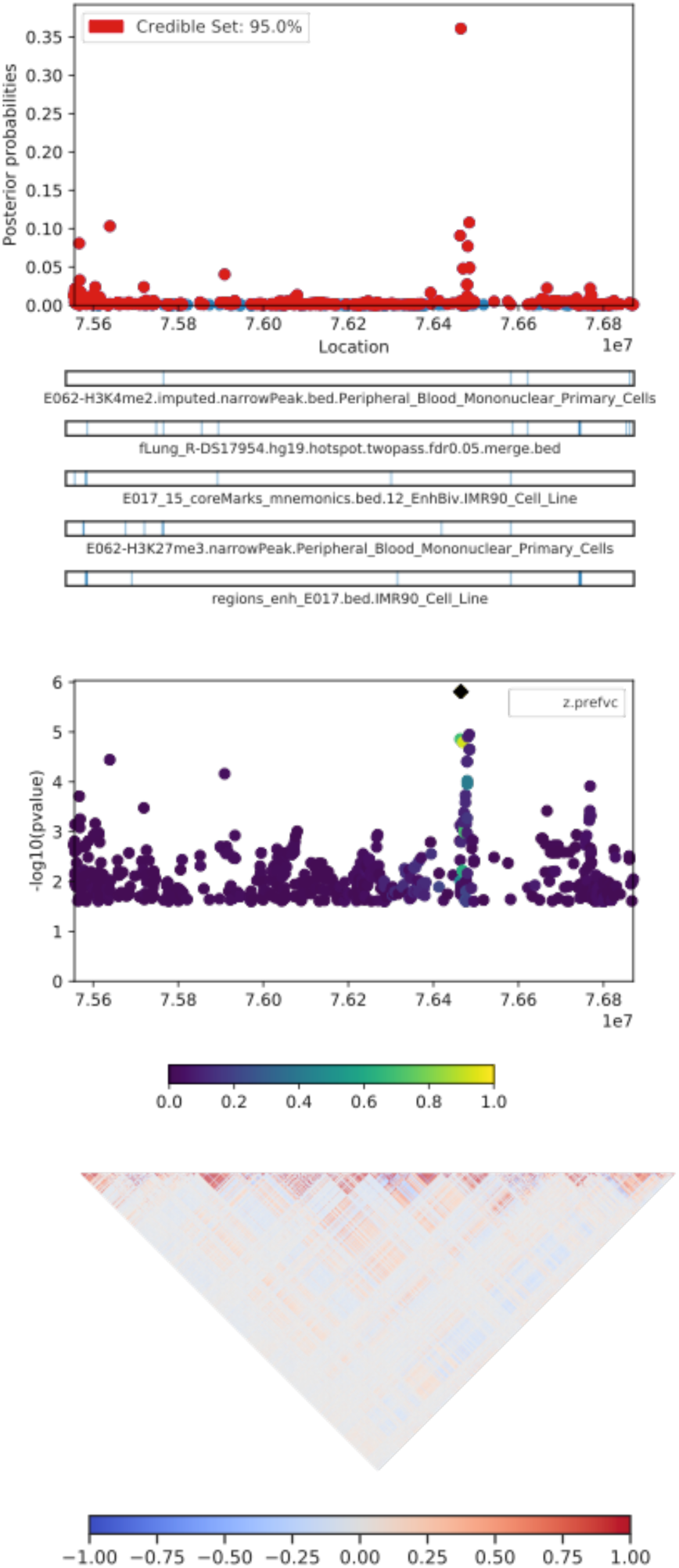
PAINTOR results for locus 2, an association on chromosome 4 with Pre-FVC. The sentinel SNP, rs10857225, corresponds with a GWAS peak that does not pass Bonferroni correction for statistical significance. The highlighted peak tags the intron of the gene *THAP6*.

**Figure 4:**
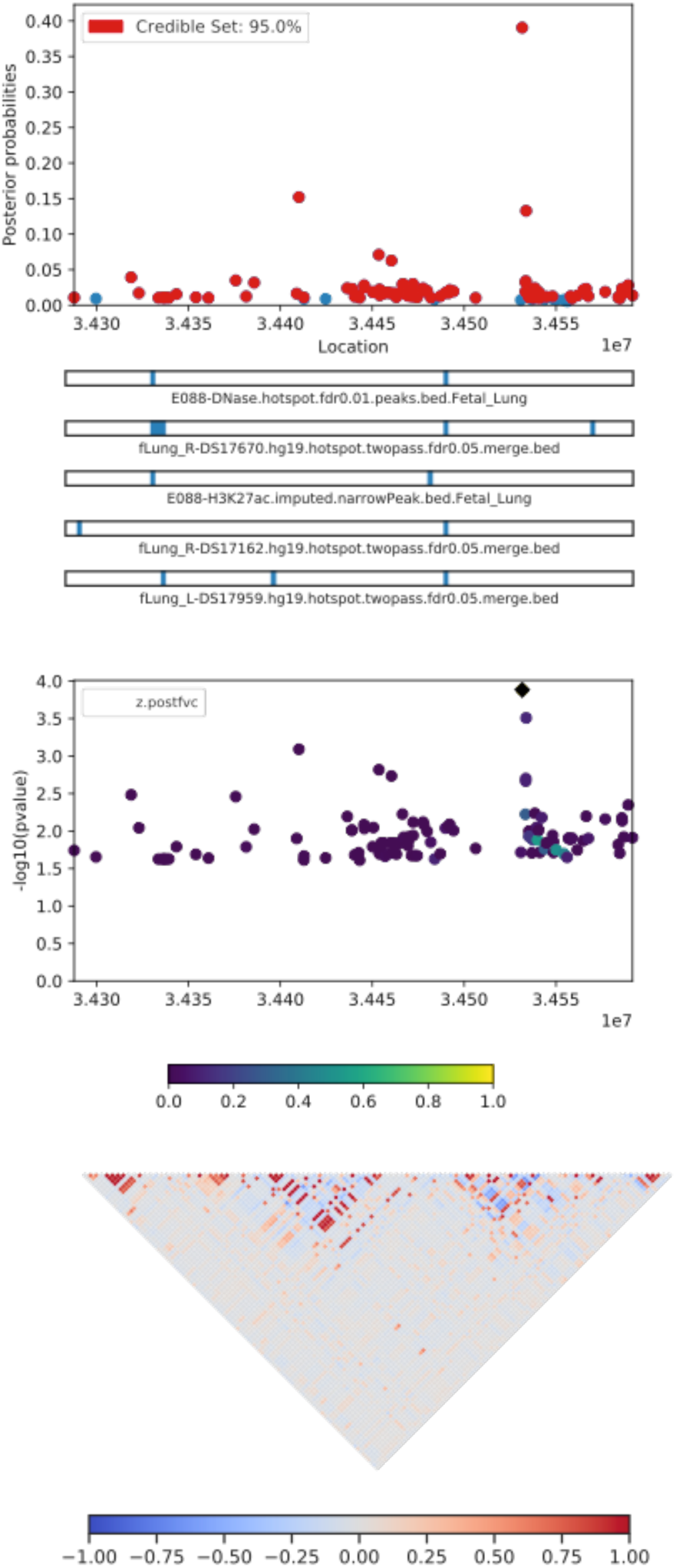
PAINTOR fine-mapping results for locus 5, corresponding to a region on chromosome 14 associated with Post-FVC. The most likely causal SNP, rs1351618, tags an intron of the gene *EGLN3*.

## Discussion

We analyzed the genetic basis of six lung function phenotypes in 1,103 African American children with and without asthma. The phenotypes consisted of three standard spirometric measures – FEV_1_, FVC, and FEV_1_/FVC – measured before and after administration of bronchodilator medication. Our GWAS identified 18 genome-wide significant loci, while our integrative genetic analysis approach that layered GWAS, admixture mapping, and functional fine-mapping identified another 5 putatively causal loci that could drive differences between pre- and post-bronchodilator lung function.

The four variants on chromosome 13 associated with Pre-FEV_1_/FVC pointed to *ATP8A2* as a candidate gene. *ATP8A2* encodes an ATPase involved in phospholipid transport and is highly expressed in brain tissue, testes, and the adrenal glands, and to a lesser degree, lung (Fagerberg et al., 2014). Mutations in *ATP8A2* have been linked to several neurological disorders (Martín-Hernández et al., 2016). Two variants on chromosome 16 that were associated with Pre-FVC point to *IRX3* as a candidate gene. *IRX3* encodes a homeobox protein crucial for neural development and its promoters previously showed a long-range interaction with the *FTO* gene. Expression levels of the *FTO* gene are known to influence BMI and are of great interest in type II diabetes and obesity research (Smemo et al., 2014). Post-FVC showed associations on two chromosomes. Notable genes near the association peak on chromosome 19 included *MAP2K2* and *ZBTB7A*, genes associated with variation in BMI, visceral adiposity, and eosinophil counts (Kichaev et al., 2019; Pulit et al., 2019; Rüeger et al., 2018); and *CHAF1A, HDGFL2, PLIN4, ANKRD24, MPND*, and *SH3GL1*, previously associated with corpuscular volume and hemoglobin concentration (Astle et al., 2016; Kichaev et al., 2019; van Rooij et al., 2017). Interestingly, the two genes closest to the association peak, *TMIGD2* and *SHD*, have not been previously associated with any traits. *TMIGD2* is involved in T-cell co-stimulation and the immune response through an interaction with *HHLA2*, suggesting that it could possibly play an immune or allergic response role in lung function (Y. Zhu et al., 2013). Among the genes within or near the chromosome 11 peak, two emerge as potentially key loci. The first is *CXCR5*, which has been linked to increased risk of childhood onset asthma (M. A. R. Ferreira et al., 2019; Johansson et al., 2019; Pividori et al., 2019) and respiratory disease (Kichaev et al., 2019), as well as related allergic and immunological conditions such as eczema, leukocyte count, rheumatoid arthritis, and Sjögren’s syndrome (M. A. Ferreira et al., 2017; Kichaev et al., 2019; Laufer et al., 2019; Lessard et al., 2013). The second is *HYOU1*, which has been associated with BMI and Post-FEV_1_/FVC (Lutz et al., 2015; Pulit et al., 2019). Post-FEV_1_/FVC was associated with two genes, *AKAP13* and *ADAMTS7P4. AKAP13* has been previously associated with numerous conditions, including interstitial lung disease and psoriasis in European individuals (Fingerlin et al., 2013; Tsoi et al., 2015) as well as weight, BMI, and cardiovascular traits such as blood pressure and hemoglobin count in multiple populations (Giri et al., 2019; Kichaev et al., 2019). *ADAMTS7P4* was previously associated with red blood cell volume (Kichaev et al., 2019). The statistically suggestive association of Post-FEV_1_ with *BTBD11* pointed to previous associations with various lung function measures, including Post-FEV_1_, Post-FEV_1_/FVC, and ΔFEV_1_ (Hardin et al., 2016; Lutz et al., 2015). These previously detected associations were based on much larger sample sizes than what was available to us: the associations with Post-FEV_1_ and Post-FEV_1_/FVC found by Lutz et al. were discovered in a population of 10,094 European and 3,260 African American smokers with chronic obstructive pulmonary disorder (COPD), while the association with ΔFEV_1_ found by Hardin et al. was based on 5,766 Europeans and 811 African Americans with COPD, suggesting that our inability to reach genome-wide significance in our sample was insufficient statistical power.

Among significant and suggestive GWAS loci, the association of Post-FVC with variants in or near *CXCR5* and *HYOU1* is the only one that replicates known lung function loci: *CXCR5* was previously associated with asthma (M. A. R. Ferreira et al., 2019; Johansson et al., 2019; Pividori et al., 2019), and *HYOU1* was previously associated with Post-FEV_1_/FVC (Lutz et al., 2015). The association with Post-FEV_1_/FVC comes from an adult COPD cohort ascertained by smoking status; in contrast, SAGE is a pediatric asthma cohort. The mechanism by which *HYOU1* affects lung function in both youth and and adults is unclear. Nevertheless, the overlap of post-bronchodilator pulmonary function measures at this locus suggests that the region encompassing *CXCR5* and *HYOU1* plays a role in lung disease among people with obstructive lung function.

Admixture mapping identified 5 genomic regions where variation in genetic ancestry was significantly associated with phenotypic variation. Locus 1 on chromosome 21 spanned the genes *ADAMTS1* and *ADAMTS5*, which encode extracellular proteases within the same protein family but with different consequences for disease. Although both genes have been linked to blood protein levels (Suhre et al., 2017), *ADAMTS1* has been associated with pre-FVC (Kichaev et al., 2019) and is expressed in arterial, adipose, and lung tissue, while *ADAMTS5* is not appreciably expressed in the lung (Supplementary Figure 16). Further fine-mapping with PAINTOR places the most likely causal SNP (rs13615) within the 3’ UTR of *ADAMTS1*, suggesting that *ADAMTS1* may be the gene that is functionally related to lung function. Interestingly, the association of *ADAMTS1* with pre-FVC (Kichaev et al., 2019) was discovered in a European sample of substantially larger size than our cohort, highlighting the ability of admixture mapping to detect associations in scenarios with low statistical power. Locus 3, spanning a region on chromosome 19 that was associated with Pre-FEV_1_/FVC, contains the genes *ZNF557* and *INSR. ZNF557* has not been previously associated with any traits, while *INSR* is the well-known insulin receptor that has been previously associated with childhood onset asthma in our own cohort (White et al., 2016), as well as blood pressure levels, triglyceride levels, HDL cholesterol levels, and hypothyroidism across multiple populations (Bentley et al., 2019; Ehret et al., 2016; Kichaev et al., 2019; Klarin et al., 2018). Post-FVC showed two distinct admixture mapping signals. The first region on chromosome 8, which we call Locus 4, includes the genes *ESRP1, INTS8, TP53INP1* and *NDUFAF6* was previously associated with type II diabetes and eosinophil counts (Kichaev et al., 2019; Mahajan et al., 2018). The second region, Locus 5, spans *EGLN3* and *SNX6*, both of which show previous associations with blood phenotypes such as blood pressure and hematocrit levels (Astle et al., 2016; Evangelou et al., 2018).

Overall, evaluation of our GWAS and admixture mapping lung function results suggests that genetics of this trait underlies some pleiotropy observed across pulmonary, hematological, cardiovascular, and obesity-related traits. Such pleiotropy has been observed in UK BioBank participants: as lung function decreases, BMI and type II diabetes incidence increases, as well as levels of eosinophils and neutrophils, both of which are common biomarkers for allergic disease (Supplementary Figure 25)(McInnes et al., 2019; Stanford Global Biobank Engine, 2020). The link between obesity and lung function is particularly interesting since obesity is a known asthma comorbidity, and lung function may play a role in obese asthma (Baffi et al., 2015)(Gruchała-Niedoszytko et al., 2015). Our findings suggest that genetically-based differences in lung function may provide a link between obesity and asthma.

It is curious that the regions identified by admixture mapping and subjected to functional fine-mapping did not overlap with the statistically significant GWAS loci. We attribute this in part to the different types of information used by each approach: GWAS analyzes how allelic variation affects a trait, while admixture mapping analyzes the phenotypic consequences of variation in genetic ancestry. By integrating GWAS summary statistics with loci identified via admixture mapping, we found that three of the admixture mapping-based loci -- *ADAMTS1, THAP6*, and *EGLN3* -- had evidence of causal effects. Each of the sentinel SNPs tagging these genes showed a notable difference in ancestral allele frequency: populations of African descent had either the highest or the lowest minor allele frequency among all global populations, likely the result of admixture mapping prioritizing loci that varied by genetic ancestry. None of these loci have been previously associated with lung traits, highlighting the strength of our integrative analysis. The association with *EGLN3* is particularly curious since it has been previously associated with a variety of traits, including heart rate response to beta-blocker therapy (Shahin et al., 2018). Short-acting beta-2 agonists such as albuterol selectively target beta-2 receptors in the lungs, while the first-generation beta-blockers taken for cardiac conditions bind to both beta-1 and beta-2 receptors, affecting the heart as well the lungs. Bronchospasm and FEV_1_ reduction are clinically significant side effects of first-generation beta-1 selective and nonselective beta-blockers for cardiac conditions. Consequently, these non-selective beta-blockers must be initiated with caution and close monitoring in patients with asthma (Christiansen & Zuraw, 2019). Beta-blockers lower blood pressure by reducing heart rate and cardiac contractility and are less effective in people with high levels of African genetic ancestry (Brewster & Seedat, 2013; Whelton et al., 2018). It has been previously observed that African Americans with asthma demonstrate lower bronchodilator drug response than European Americans (Blake et al., 2008), suggesting a possible pharmacological interaction between beta-2 receptors and African ancestry. Furthermore, *EGLN3* is strongly expressed in cardiac tissue, suggesting that *EGLN3* could possibly influence Post-FVC through cardiac phenotypes (Supplementary Figure 24). Further functional studies are required to elucidate the role of *EGLN3* on lung function and bronchodilator drug response.

Our integrative analysis approach leverages available functional annotations and genetic ancestry estimates in the absence of molecular data to yield some promise for discovery of novel loci. Our study is limited to three tiers – genotypes, genetic ancestry, and functional annotations – and makes use of gene expression results from GTEx v8. However, it does not directly incorporate any transcriptomic, metabolomic, proteomic, or methylomic information. As large multi-omic datasets from NHLBI TOPMed, UK Biobank, and the NIH Million Veterans Program become available, the need for integrative genomic approaches to studying complex diseases will increase. Future multi-omic models of complex diseases, including obstructive lung function disorders, may deliver on the promise of precision medicine and provide actionable clinical translation of biomedical and pharmacogenomic insights into novel therapies.

## Acknowledgements

This work was supported in part by the Sandler Family Foundation, the American Asthma Foundation, the RWJF Amos Medical Faculty Development Program, the Harry Wm. and Diana V. Hind Distinguished Professor in Pharmaceutical Sciences II, the National Heart, Lung, and Blood Institute (NHLBI) grants R01HL117004, R01HL128439, R01HL135156, X01HL134589, R01HL141992, R01HL104608, R01HL141845, and U01HL138626, the National Human Genome Research Institute (NHGRI) grants U01HG007419 and U01HG009080, the National Institute of Environmental Health Sciences grants R01ES015794, R21ES24844, the Eunice Kennedy Shriver National Institute of Child Health and Human Development (NICHD) grant R01HD085993, the National Institute on Minority Health and Health Disparities (NIMHD) grants P60MD006902, R01MD010443, RL5GM118984, R56MD013312, and R56MD013312, and the Tobacco-Related Disease Research Program under Award Numbers 24RT-0025 and 27IR-0030. Research reported in this article was funded by the National Institutes of Health Common Fund and Office of Scientific Workforce Diversity under three linked awards RL5GM118984, TL4GM118986, 1UL1GM118985 administered by the National Institute of General Medical Sciences (NIGMS).

The authors wish to acknowledge the following SAGE co-investigators for subject recruitment, sample processing and quality control: Luisa N. Borrell, DDS, PhD, Emerita Brigino-Buenaventura, MD, Adam Davis, MA, MPH, Michael A. LeNoir, MD, Kelley Meade, MD, Fred Lurmann, MS and Harold J. Farber, MD, MSPH. The authors also wish to thank the staff and participants who contributed to the SAGE study.

This manuscript uses results and visualization provided by the Stanford Global Biobank Engine. The authors would like to thank the Rivas Lab for making the resource available.

PCG was additionally funded by NHGRI training grant T32 HG000044 to the Department of Genetics at Stanford University.

KLK was additionally supported by NHLBI grant supplement R01HL135156-S1, the UCSF Bakar Computational Health Sciences Institute, the Gordon and Betty Moore Foundation grant GBMF3834, and the Alfred P. Sloan Foundation grant 2013-10-27 to UC Berkeley through the Moore-Sloan Data Sciences Environment initiative at the Berkeley Institute for Data Science (BIDS). The logistical space, technical support, administrative assistance, and indefatigable good humor of the members and staff at BIDS are gratefully acknowledged.

EYL, LSB, and AKL were supported by a National Research Service Award grant T32GM007546 from the NIGMS. MGC was additionally supported by NIH MARC U-STAR grant T34GM008574 at San Francisco State University. MJW was additionally supported by NHLBI grant supplement R01HL117004-S1, an NIGMS Institutional Research and Academic Career Development Award K12GM081266, and an NHLBI Research Career Development Award K01HL140218.

## Author Contributions

PCG, KLK, and MJW designed the study. SSO, SS, CE, and EGB recruited study subjects and generated the data. ACYM, JRE, DU, and SH cleaned and organized the data and provided analytic support. PCG, KLK, EYL, OR, MGC, and MJW performed the analysis. AKL and LSB provided clinical pharmacological expertise for interpretation of results. EGB and BEH funded the study. EGB supervised all recruitment. All authors contributed to manuscript writing and editing.

## Supplementary Tables and Figures

**Supplementary Table 1:** Counts of individuals and variants retained and removed during quality control, including filtering flags from PLINK 1.9. Initial indicates the starting number of individuals and variants in the unfiltered datasets. The subsequent row names refer to the plink command and threshold used for each step: (1) Individuals whose DNA was obtained from saliva rather than blood were removed; (2) Variants with genotyping efficiency below 95% were purged; (3) Variants with minor allele frequencies below 1% were excluded; (4) Individuals with genotyping efficiency below 95% were removed. (5) Variants that deviate from Hardy-Weinberg equilibrium with p-value < 0.0001 were purged.

|  | Individuals (*n*) |  | Genotyped (*n*) |  | Imputed (*n*) |  |
| --- | --- | --- | --- | --- | --- | --- |
|  | *retained* | *removed* | *retained* | *removed* | *retained* | *removed* |
| Initial | 1985 | - | 772,703 | - | 47,101,126 | - |
| (1) remove saliva | 1949 | 36 | 772,703 | 0 | 47,101,126 | 0 |
| (2) --geno 0.05 | 1949 | 0 | 744,492 | 28,211 | 47,101,126 | 0 |
| (3) --maf 0.01 | 1949 | 0 | 664,072 | 80,420 | 16,005,708 | 31,095,418 |
| (4) --mind 0.05 | 1948 | 1 | 664,072 | 0 | 16,005,708 | 0 |
| (5) --hwe 0.0001 | 1948 | 0 | 660,802 | 3,270 | 15,954,804 | 50,904 |
| **Final** | **1948** | **37** | **660,802** | **31,561** | **15,954,804** | **31,146,322** |

**Supplementary Table 2:** Genomic inflation factors (λ), effective numbers of tests (M_eff_), and both significance and suggestive thresholds are presented for the GWAS of each phenotype. The genomic inflation values were calculated in R. The number M_eff_ was estimated in R using the autocorrelation method from CODA. The significance threshold was calculated as 1/ M_eff_, and the suggestive threshold was determined with 0.5/ M_eff_.

| GWAS Trait | $\lambda$ | Effective Tests | Significance Threshold | Suggestive Threshold |
| --- | --- | --- | --- | --- |
| Post-FEV<sub>1</sub>/FVC | 0.9776 | 503667.1 | 9.93x 10<sup>-8</sup> | 1.99 x 10<sup>-7</sup> |
| Post-FEV<sub>1</sub> | 0.9791 | 512041 | 9.76 x 10<sup>-8</sup> | 1.95 x 10<sup>-7</sup> |
| Post-FVC | 0.9732 | 488818.5 | 1.02 x 10<sup>-7</sup> | 2.05 x 10<sup>-7</sup> |
| Pre-FEV<sub>1</sub>/FVC | 0.9885 | 499624.2 | 1.00 x 10<sup>-7</sup> | 2.00 x 10<sup>-7</sup> |
| Pre-FEV<sub>1</sub> | 0.9870 | 507975.4 | 9.84 x 10<sup>-8</sup> | 1.97 x 10<sup>-7</sup> |
| Pre-FVC | 0.9893 | 502450.9 | 9.95 x 10<sup>-8</sup> | 1.99 x 10<sup>-7</sup> |
| **average** | **0.9824** | **502429.5** | **9.95 x 10<sup>-8</sup>** | **1.99 x 10<sup>-7</sup>** |

**Supplementary Figure 1:**
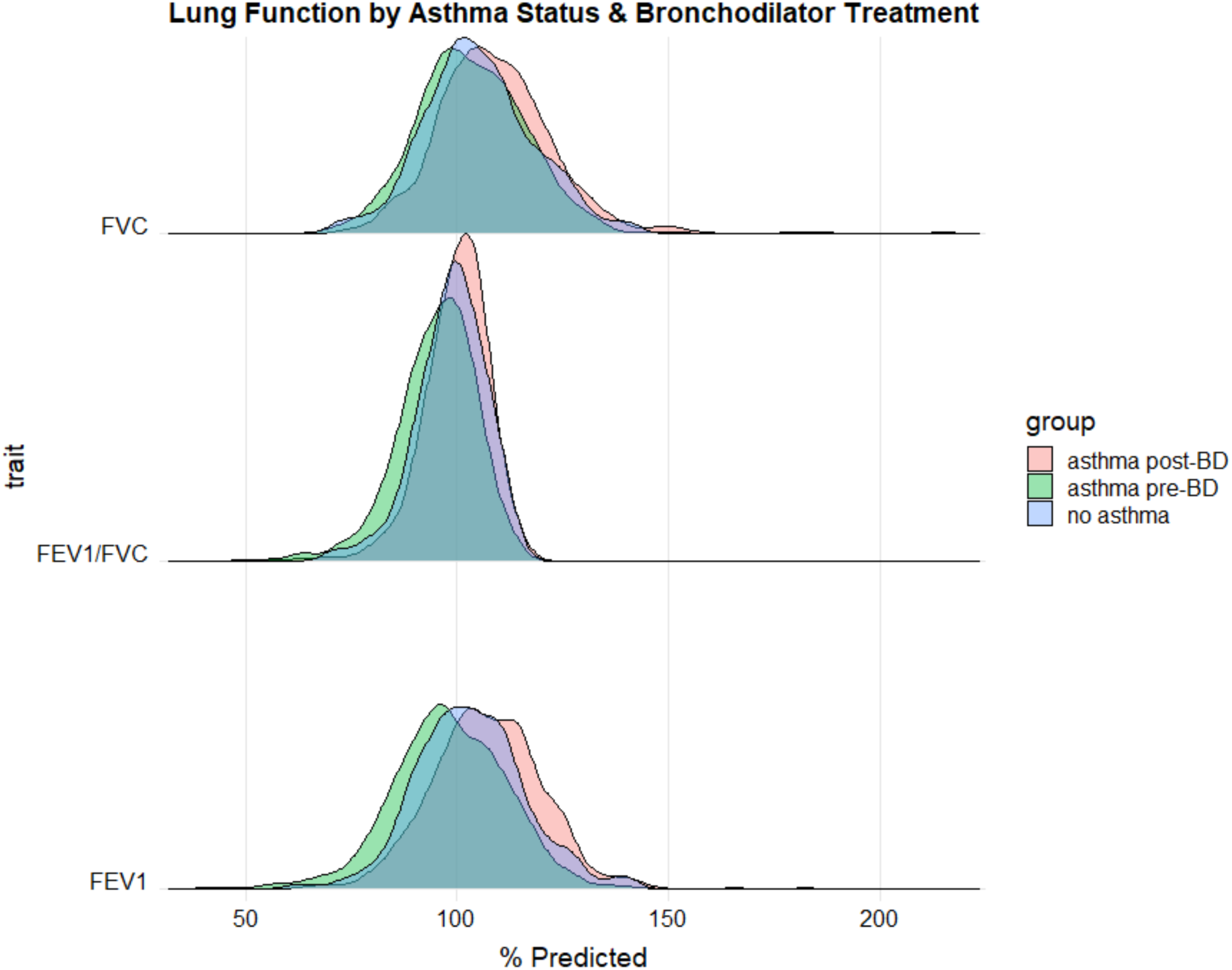
Distributions of lung function measures stratified by asthma case status and bronchodilator treatment. Lung function values are normalized against predictions given by the Hankinson equations (% Predicted). Asthma controls (blue) received no bronchodilator medication. Asthma cases are separated by lung function values measured pre-bronchodilator (pre-BD, green) and post-bronchodilator (post-BD, red) administration.

**Supplementary Figure 2:**
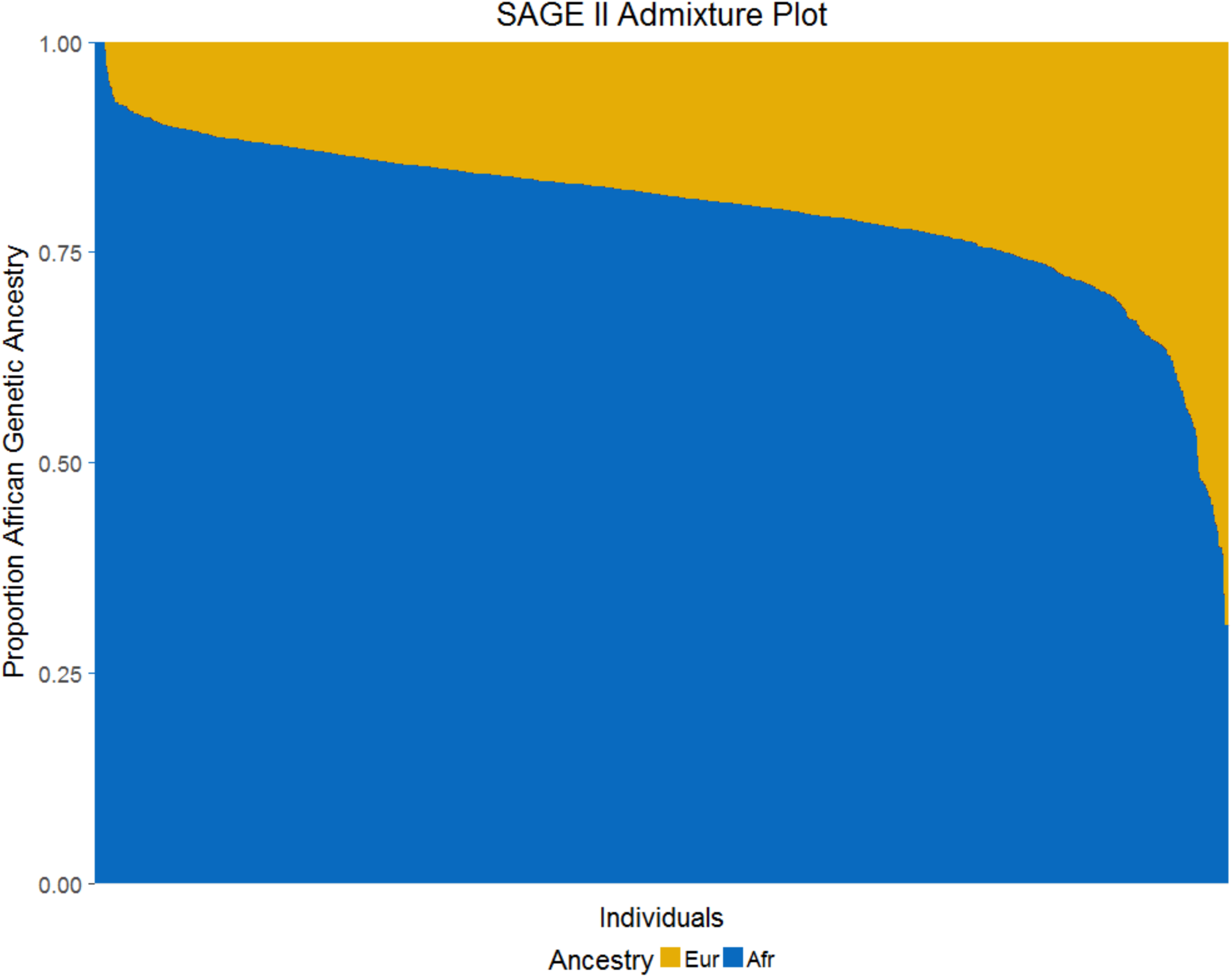
A plot of global admixture proportions for SAGE subjects, ordered in descending proportion of African ancestry. In SAGE, the mean genetic ancestry proportion of African origin is 80.2%.

**Supplementary Figure 3:**
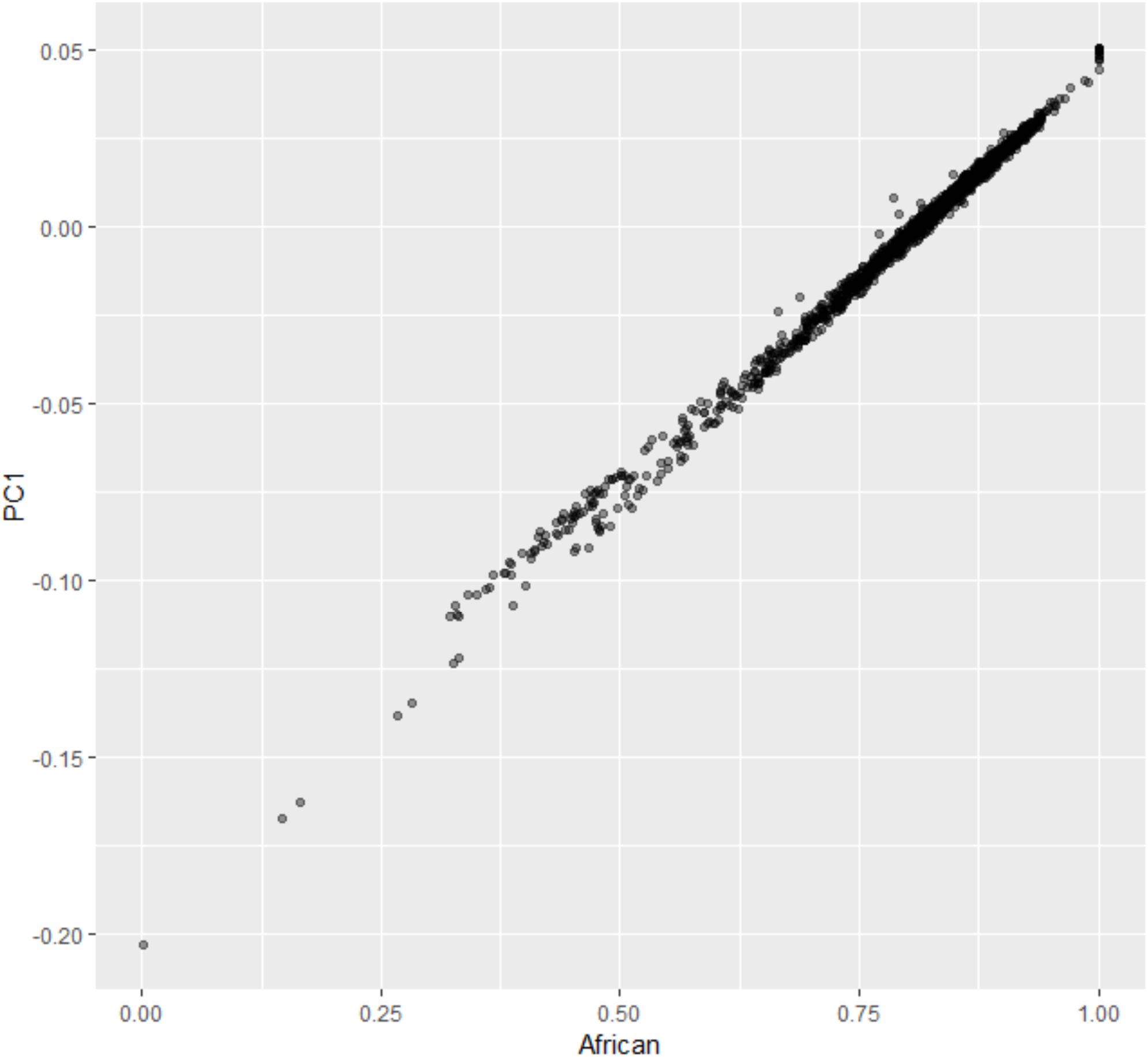
A plot of the first genotype principal component (PC1) versus the proportion of global African genetic ancestry (African) shows that the two covariates are essentially exchangeable (*R*^2^ = 0.984).

**Supplementary Figure 4:**
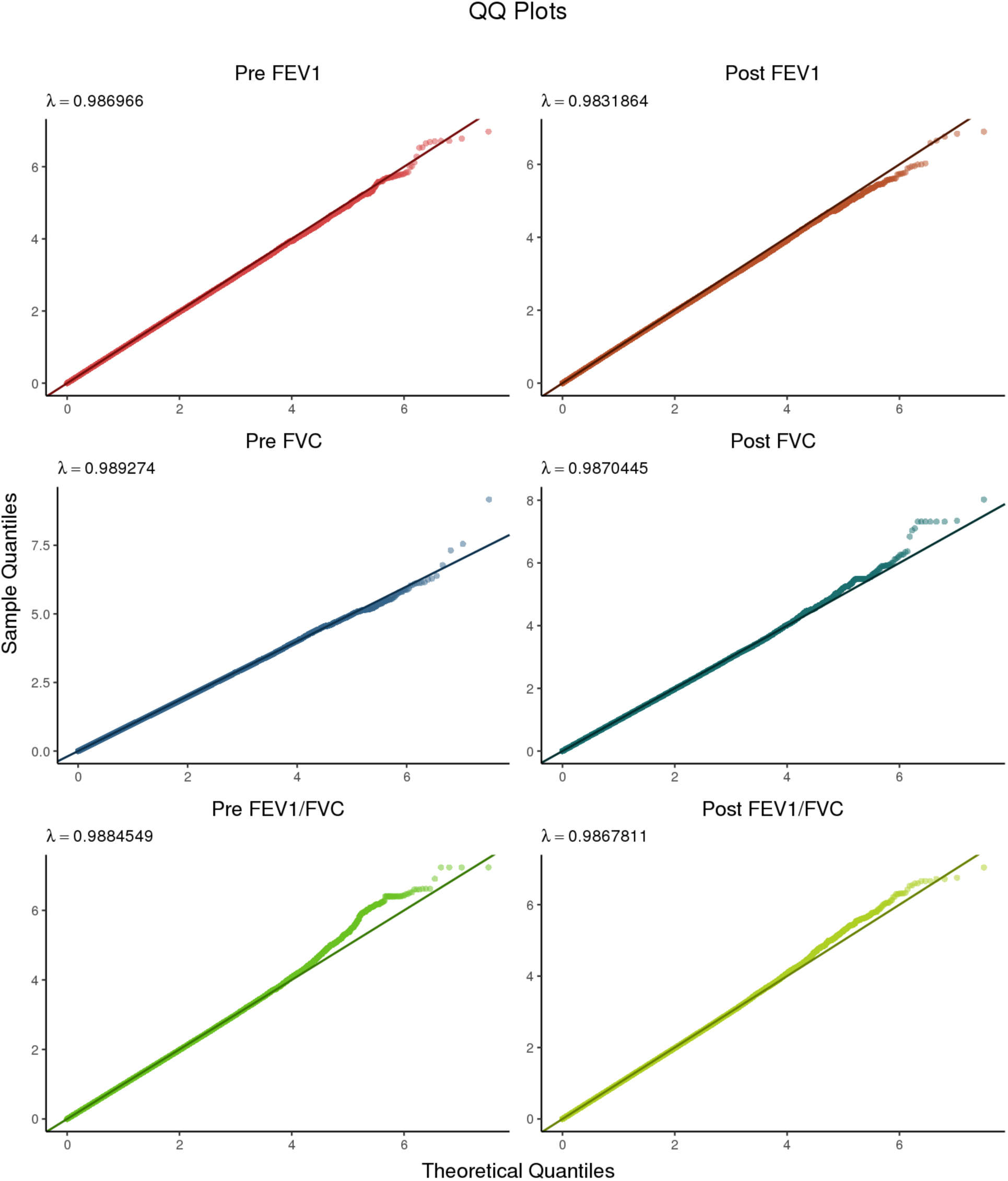
Quantile-quantile plots for all six phenotypes subjected to GWAS. The left column contains Pre-BD phenotypes, while the right column contains the corresponding Post-BD phenotypes.

**Supplementary Figure 5:**
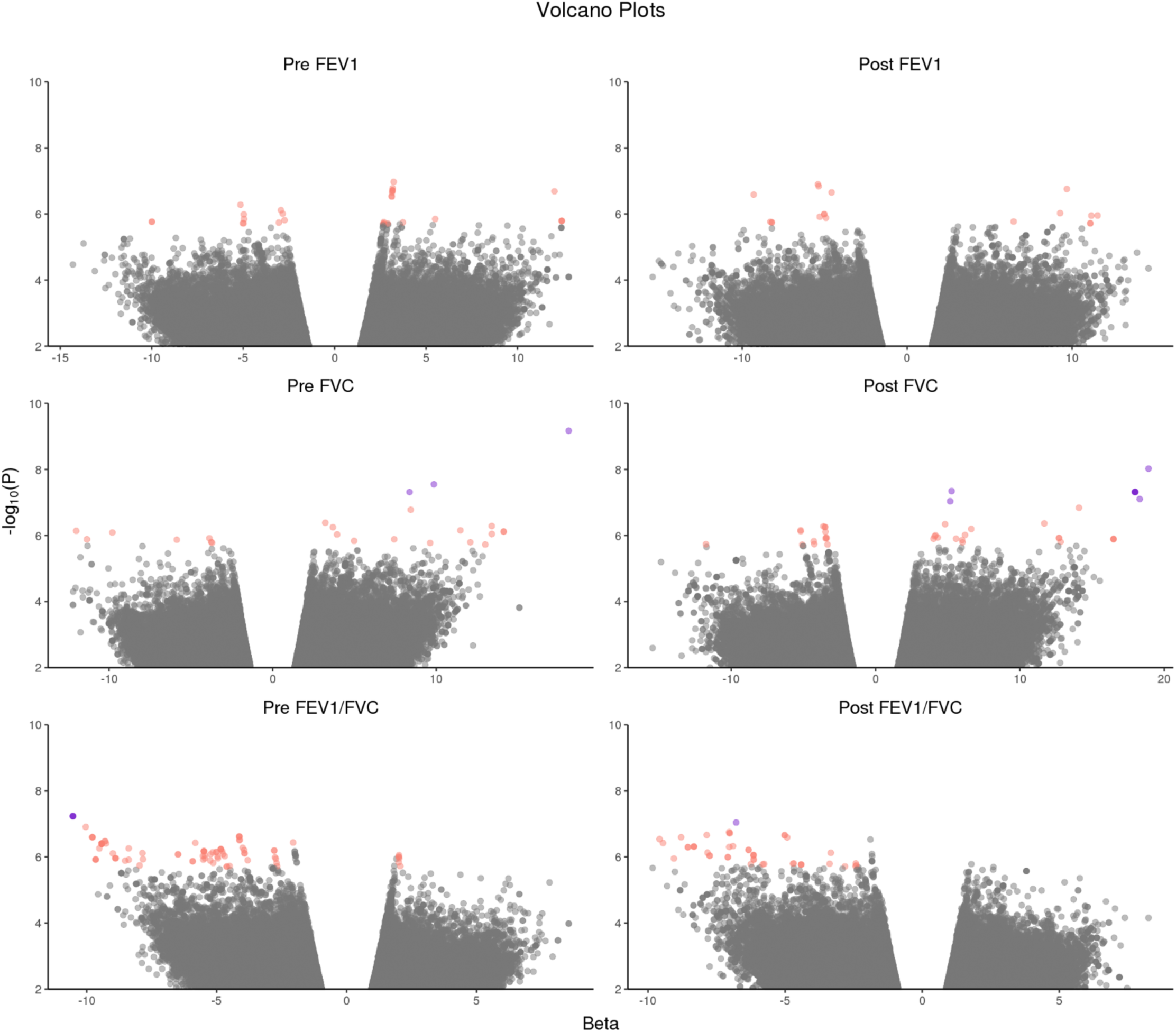
Volcano plots showing the relationship between GWAS *p*-values and effect sizes β. As in Supplementary Figure 4, pre-BD traits are on the left, while post-BD traits are on the right. Variants meeting the CODA-adjusted suggestive threshold (1.99 × 10^−6^) are highlighted in salmon pink; variants with a *p*-value ≤ 1 × 10^−7^ and |β| ≥ 2 are highlighted in purple. A positive value of β indicates that the minor allele is associated with higher lung function, while β < 0 indicates that the minor allele tracks with lower lung function compared to the major allele. Non-significant variants are shown in grey. Variants with a *p*-value < 0.05 were deemed uninformative and therefore not plotted.

**Supplementary Table 3:** Suggestive associations with Pre-FEV_1_

| Phenotype | Chr | SNP | Position (bp) | A1 | A2 | MAF | $\beta$ | Std Err | P-value |
| --- | --- | --- | --- | --- | --- | --- | --- | --- | --- |
| Pre-FEV<sub>1</sub> | 1 | 1:158003840 | 158003840 | C | G | 0.012 | 12.4180 | 2.59 | 1.61 x 10<sup>-6</sup> |
| Pre-FEV<sub>1</sub> | 1 | 1:158004676 | 158004676 | C | T | 0.011 | 12.4151 | 2.59 | 1.62 x 10<sup>-6</sup> |
| Pre-FEV<sub>1</sub> | 2 | 2:34321239 | 34321239 | C | T | 0.293 | -3.0486 | 0.639 | 1.82 x 10<sup>-6</sup> |
| Pre-FEV<sub>1</sub> | 6 | 6:12318099 | 12318099 | A | G | 0.081 | -4.9768 | 1.05 | 1.92 x 10<sup>-6</sup> |
| Pre-FEV<sub>1</sub> | 10 | 10:109510095 | 109510095 | T | G | 0.161 | 3.7327 | 0.782 | 1.79 x 10<sup>-6</sup> |
| Pre-FEV<sub>1</sub> | 10 | 10:109519015 | 109519015 | A | G | 0.405 | -2.7360 | 0.569 | 1.54 x 10<sup>-6</sup> |
| Pre-FEV<sub>1</sub> | 10 | 10:109531399 | 109531399 | C | T | 0.330 | 3.1622 | 0.607 | 1.93 x 10<sup>-7</sup> |
| Pre-FEV<sub>1</sub> | 10 | 10:109531773 | 109531773 | A | G | 0.331 | 3.1622 | 0.607 | 1.93 x 10<sup>-7</sup> |
| Pre-FEV<sub>1</sub> | 10 | 10:109532399 | 109532399 | G | A | 0.331 | 3.1409 | 0.607 | 2.24 x 10<sup>-7</sup> |
| Pre-FEV<sub>1</sub> | 10 | 10:109532529 | 109532529 | G | T | 0.331 | 3.1755 | 0.607 | 1.66 x 10<sup>-7</sup> |
| Pre-FEV<sub>1</sub> | 10 | 10:109533077 | 109533077 | C | T | 0.331 | 3.1174 | 0.608 | 2.89 x 10<sup>-7</sup> |
| Pre-FEV<sub>1</sub> | 10 | 10:109538187 | 109538187 | A | C | 0.381 | -2.8430 | 0.580 | 9.71 x 10<sup>-7</sup> |
| Pre-FEV<sub>1</sub> | 10 | 10:109538400 | 109538400 | T | C | 0.328 | 3.1254 | 0.610 | 2.98 x 10<sup>-7</sup> |
| Pre-FEV<sub>1</sub> | 10 | 10:109542795 | 109542795 | C | T | 0.318 | 2.9032 | 0.610 | 1.94 x 10<sup>-6</sup> |
| Pre-FEV<sub>1</sub> | 10 | 10:109543483 | 109543483 | G | A | 0.328 | 3.1689 | 0.609 | 1.98 x 10<sup>-7</sup> |
| Pre-FEV<sub>1</sub> | 12 | 12:125410562 | 125410562 | T | G | 0.336 | -2.9360 | 0.594 | 7.68 x 10<sup>-7</sup> |
| Pre-FEV<sub>1</sub> | 14 | 14:20641690 | 20641690 | C | T | 0.410 | 2.6744 | 0.563 | 1.99 x 10<sup>-6</sup> |
| Pre-FEV<sub>1</sub> | 14 | 14:20641707 | 20641707 | T | A | 0.410 | 2.6744 | 0.563 | 1.99 x 10<sup>-6</sup> |
| Pre-FEV<sub>1</sub> | 14 | 14:20641788 | 20641788 | A | T | 0.410 | 2.6795 | 0.563 | 1.93 x 10<sup>-6</sup> |
| Pre-FEV<sub>1</sub> | 14 | 14:20642034 | 20642034 | A | T | 0.410 | 2.6878 | 0.562 | 1.76 x 10<sup>-6</sup> |
| Pre-FEV<sub>1</sub> | 17 | 17:1307087 | 1307087 | A | G | 0.087 | -5.0185 | 1.05 | 1.90 x 10<sup>-6</sup> |
| Pre-FEV<sub>1</sub> | 17 | 17:10856828 | 10856828 | G | A | 0.065 | 5.4992 | 1.14 | 1.42 x 10<sup>-6</sup> |
| Pre-FEV<sub>1</sub> | 17 | 17:66947338 | 66947338 | G | A | 0.017 | -9.9925 | 2.09 | 1.72 x 10<sup>-6</sup> |
| Pre-FEV<sub>1</sub> | 17 | 17:66947385 | 66947385 | G | A | 0.017 | -9.9925 | 2.09 | 1.72 x 10<sup>-6</sup> |
| Pre-FEV<sub>1</sub> | 19 | 19:45780813 | 45780813 | G | A | 0.072 | -4.9503 | 1.03 | 1.42 x 10<sup>-6</sup> |
| Pre-FEV<sub>1</sub> | 19 | 19:45782169 | 45782169 | T | C | 0.073 | -4.9651 | 1.02 | 1.03 x 10<sup>-6</sup> |
| Pre-FEV<sub>1</sub> | 19 | 19:45782594 | 45782594 | G | A | 0.073 | -5.1489 | 1.03 | 5.24 x 10<sup>-7</sup> |
| Pre-FEV<sub>1</sub> | 20 | 20:42117790 | 42117790 | T | C | 0.016 | 12.0182 | 2.31 | 2.05 x 10<sup>-7</sup> |

**Supplementary Table 4:** Suggestive associations with Pre-FVC

| Phenotype | Chr | SNP | Position (bp) | A1 | A2 | MAF | $\beta$ | Std Err | P-value |
| --- | --- | --- | --- | --- | --- | --- | --- | --- | --- |
| Pre-FVC | 3 | 3:136065565 | 136065565 | G | A | 0.137 | 3.9445 | 0.804 | $9.28 \times 10^{-7}$ |
| Pre-FVC | 3 | 3:136073920 | 136073920 | T | C | 0.165 | 3.6839 | 0.736 | $5.62 \times 10^{-7}$ |
| Pre-FVC | 3 | 3:140836152 | 140836152 | A | G | 0.011 | 12.0836 | 2.52 | $1.60 \times 10^{-6}$ |
| Pre-FVC | 3 | 3:172005706 | 172005706 | G | T | 0.026 | 8.4445 | 1.61 | $1.67 \times 10^{-7}$ |
| Pre-FVC | 6 | 6:45961727 | 45961727 | T | G | 0.017 | 9.6273 | 2.01 | $1.68 \times 10^{-6}$ |
| Pre-FVC | 6 | 6:162713288 | 162713288 | A | G | 0.014 | 11.4754 | 2.31 | $6.98 \times 10^{-7}$ |
| Pre-FVC | 11 | 11:86410675 | 86410675 | A | G | 0.010 | 13.3956 | 2.73 | $8.98 \times 10^{-7}$ |
| Pre-FVC | 14 | 14:51754057 | 51754057 | T | C | 0.216 | 3.2204 | 0.636 | $4.11 \times 10^{-7}$ |
| Pre-FVC | 15 | 15:68708723 | 68708723 | T | C | 0.142 | -3.7504 | 0.781 | $1.57 \times 10^{-6}$ |
| Pre-FVC | 16 | 16:54327610 | 54327610 | A | G | 0.034 | 8.3689 | 1.53 | $4.85 \times 10^{-8}$ |
| Pre-FVC | 16 | 16:54327903 | 54327903 | G | A | 0.023 | 9.8543 | 1.78 | $2.83 \times 10^{-8}$ |
| Pre-FVC | 19 | 19:7732700 | 7732700 | C | T | 0.030 | 7.4308 | 1.54 | $1.30 \times 10^{-6}$ |
| Pre-FVC | 19 | 19:36353880 | 36353880 | A | G | 0.121 | -3.8618 | 0.796 | $1.21 \times 10^{-6}$ |
| Pre-FVC | 19 | 19:36372775 | 36372775 | T | C | 0.129 | -3.7039 | 0.773 | $1.67 \times 10^{-6}$ |
| Pre-FVC | 20 | 20:22900228 | 22900228 | G | A | 0.010 | 18.0953 | 2.93 | $6.77 \times 10^{-7}$ |
| Pre-FVC | 20 | 20:22960678 | 22960678 | A | G | 0.012 | 13.3932 | 2.67 | $5.17 \times 10^{-7}$ |
| Pre-FVC | 20 | 20:49240699 | 49240699 | G | A | 0.070 | 4.9814 | 1.03 | $1.46 \times 10^{-6}$ |
| Pre-FVC | 20 | 20:58734941 | 58734941 | A | T | 0.010 | 14.1234 | 2.86 | $7.61 \times 10^{-7}$ |
| Pre-FVC | 20 | 20:58735391 | 58735391 | C | G | 0.010 | 14.1234 | 2.86 | $7.61 \times 10^{-7}$ |
| Pre-FVC | 21 | 21:25030141 | 25030141 | G | T | 0.015 | -12.0039 | 2.42 | $7.24 \times 10^{-7}$ |
| Pre-FVC | 21 | 21:25045811 | 25045811 | A | G | 0.016 | -11.3463 | 2.34 | $1.30 \times 10^{-6}$ |
| Pre-FVC | 21 | 21:26032912 | 26032912 | C | G | 0.011 | 12.9973 | 2.73 | $1.87 \times 10^{-6}$ |
| Pre-FVC | 22 | 22:35972126 | 35972126 | G | A | 0.052 | -5.8582 | 1.21 | $1.34 \times 10^{-6}$ |

**Supplementary Table 5:** Suggestive associations with Pre-FEV_1_/FVC.

| Phenotype | Chr | SNP | Position (bp) | A1 | A2 | MAF | $\beta$ | Std Err | P-value |
| --- | --- | --- | --- | --- | --- | --- | --- | --- | --- |
| Pre-FEV<sub>1</sub>/FVC | 2 | 2:37977506 | 37977506 | C | T | 0.035 | -4.9670 | 1.01 | 9.81 x 10<sup>-7</sup> |
| Pre-FEV<sub>1</sub>/FVC | 2 | 2:236936348 | 236936348 | C | G | 0.469 | 1.8358 | 0.385 | 1.85 x 10<sup>-6</sup> |
| Pre-FEV<sub>1</sub>/FVC | 3 | 3:31401513 | 31401513 | A | T | 0.020 | -6.4871 | 1.32 | 8.34 x 10<sup>-7</sup> |
| Pre-FEV<sub>1</sub>/FVC | 3 | 3:31401514 | 31401514 | C | T | 0.020 | -6.4871 | 1.32 | 8.34 x 10<sup>-7</sup> |
| Pre-FEV<sub>1</sub>/FVC | 4 | 4:96879482 | 96879482 | C | T | 0.019 | -7.8595 | 1.59 | 7.62 x 10<sup>-7</sup> |
| Pre-FEV<sub>1</sub>/FVC | 4 | 4:96913583 | 96913583 | C | T | 0.011 | -9.2573 | 1.82 | 3.89 x 10<sup>-7</sup> |
| Pre-FEV<sub>1</sub>/FVC | 5 | 5:49943626 | 49943626 | A | G | 0.465 | -1.9196 | 0.398 | 1.44 x 10<sup>-6</sup> |
| Pre-FEV<sub>1</sub>/FVC | 5 | 5:49943944 | 49943944 | G | A | 0.465 | -1.9191 | 0.399 | 1.48 x 10<sup>-6</sup> |
| Pre-FEV<sub>1</sub>/FVC | 5 | 5:49980285 | 49980285 | G | C | 0.476 | -1.9524 | 0.398 | 9.30 x 10<sup>-7</sup> |
| Pre-FEV<sub>1</sub>/FVC | 5 | 5:50028474 | 50028474 | A | G | 0.481 | -1.9960 | 0.402 | 6.79 x 10<sup>-7</sup> |
| Pre-FEV<sub>1</sub>/FVC | 5 | 5:50033254 | 50033254 | A | C | 0.481 | -1.9960 | 0.402 | 6.79 x 10<sup>-7</sup> |
| Pre-FEV<sub>1</sub>/FVC | 5 | 5:50068245 | 50068245 | T | A | 0.481 | -1.9881 | 0.403 | 8.07 x 10<sup>-7</sup> |
| Pre-FEV<sub>1</sub>/FVC | 5 | 5:50089919 | 50089919 | A | G | 0.482 | -1.9828 | 0.402 | 8.18 x 10<sup>-7</sup> |
| Pre-FEV<sub>1</sub>/FVC | 5 | 5:50140456 | 50140456 | A | G | 0.471 | -1.9802 | 0.403 | 9.11 x 10<sup>-7</sup> |
| Pre-FEV<sub>1</sub>/FVC | 5 | 5:50140616 | 50140616 | C | T | 0.477 | -2.0583 | 0.405 | 3.66 x 10<sup>-7</sup> |
| Pre-FEV<sub>1</sub>/FVC | 7 | 7:23773818 | 23773818 | C | T | 0.408 | 1.9154 | 0.394 | 1.14 x 10<sup>-6</sup> |
| Pre-FEV<sub>1</sub>/FVC | 7 | 7:23819271 | 23819271 | G | T | 0.288 | 2.0651 | 0.433 | 1.90 x 10<sup>-6</sup> |
| Pre-FEV<sub>1</sub>/FVC | 7 | 7:23819738 | 23819738 | C | G | 0.343 | 2.0170 | 0.410 | 8.82 x 10<sup>-7</sup> |
| Pre-FEV<sub>1</sub>/FVC | 7 | 7:23820068 | 23820068 | A | T | 0.343 | 2.0105 | 0.411 | 9.95 x 10<sup>-7</sup> |
| Pre-FEV<sub>1</sub>/FVC | 7 | 7:23821784 | 23821784 | C | A | 0.343 | 2.0105 | 0.411 | 9.95 x 10<sup>-7</sup> |
| Pre-FEV<sub>1</sub>/FVC | 7 | 7:23823355 | 23823355 | G | A | 0.329 | 2.0050 | 0.416 | 1.44 x 10<sup>-6</sup> |
| Pre-FEV<sub>1</sub>/FVC | 7 | 7:23824013 | 23824013 | G | C | 0.330 | 2.0254 | 0.416 | 1.10 x 10<sup>-6</sup> |
| Pre-FEV<sub>1</sub>/FVC | 7 | 7:52038019 | 52038019 | G | A | 0.011 | -9.6564 | 1.99 | 1.20 x 10<sup>-6</sup> |
| Pre-FEV<sub>1</sub>/FVC | 7 | 7:52041108 | 52041108 | C | G | 0.011 | -9.6564 | 1.99 | 1.20 x 10<sup>-6</sup> |
| Pre-FEV<sub>1</sub>/FVC | 7 | 7:52042215 | 52042215 | A | C | 0.011 | -9.6564 | 1.99 | 1.20 x 10<sup>-6</sup> |
| Pre-FEV<sub>1</sub>/FVC | 10 | 10:4647237 | 4647237 | C | T | 0.015 | -7.8359 | 1.61 | 1.17 x 10<sup>-6</sup> |
| Pre-FEV<sub>1</sub>/FVC | 10 | 10:4706126 | 4706126 | A | G | 0.040 | -4.6870 | 0.956 | 9.48 x 10<sup>-7</sup> |
| Pre-FEV<sub>1</sub>/FVC | 10 | 10:24126147 | 24126147 | T | A | 0.010 | -8.3860 | 1.73 | 1.22 x 10<sup>-6</sup> |
| Pre-FEV<sub>1</sub>/FVC | 10 | 10:69368950 | 69368950 | C | T | 0.037 | -4.6094 | 0.969 | 1.95 x 10<sup>-6</sup> |
| Pre-FEV<sub>1</sub>/FVC | 10 | 10:69373810 | 69373810 | A | G | 0.037 | -4.6094 | 0.969 | 1.95 x 10<sup>-6</sup> |
| Pre-FEV<sub>1</sub>/FVC | 10 | 10:69434683 | 69434683 | G | A | 0.038 | -4.7908 | 0.964 | 6.73 x 10<sup>-7</sup> |
| Pre-FEV<sub>1</sub>/FVC | 10 | 10:69471854 | 69471854 | A | G | 0.038 | -4.8378 | 0.968 | 5.81 x 10<sup>-7</sup> |
| Pre-FEV<sub>1</sub>/FVC | 10 | 10:69493149 | 69493149 | C | T | 0.038 | -4.8378 | 0.968 | 5.81 x 10<sup>-7</sup> |
| Pre-FEV<sub>1</sub>/FVC | 10 | 10:69512750 | 69512750 | G | A | 0.039 | -4.7482 | 0.963 | 8.23 x 10<sup>-7</sup> |
| Pre-FEV<sub>1</sub>/FVC | 10 | 10:69543254 | 69543254 | A | G | 0.033 | -5.1457 | 1.04 | 7.20 x 10<sup>-7</sup> |
| Pre-FEV<sub>1</sub>/FVC | 10 | 10:69583770 | 69583770 | T | C | 0.030 | -5.2467 | 1.08 | 1.28 x 10<sup>-6</sup> |
| Pre-FEV<sub>1</sub>/FVC | 10 | 10:69600576 | 69600576 | T | C | 0.051 | -4.1251 | 0.806 | 3.10 x 10<sup>-7</sup> |
| Pre-FEV<sub>1</sub>/FVC | 10 | 10:69607048 | 69607048 | C | T | 0.052 | -4.1152 | 0.803 | 3.02 x 10<sup>-7</sup> |
| Pre-FEV<sub>1</sub>/FVC | 10 | 10:69622074 | 69622074 | A | T | 0.052 | -3.9272 | 0.795 | 7.74 x 10<sup>-7</sup> |
| Pre-FEV<sub>1</sub>/FVC | 10 | 10:69622076 | 69622076 | G | A | 0.052 | -3.9272 | 0.795 | 7.74 x 10<sup>-7</sup> |
| Pre-FEV<sub>1</sub>/FVC | 10 | 10:69629302 | 69629302 | A | G | 0.028 | -5.4903 | 1.10 | 6.68 x 10<sup>-7</sup> |
| Pre-FEV<sub>1</sub>/FVC | 10 | 10:69629812 | 69629812 | T | G | 0.028 | -5.4903 | 1.10 | 6.68 x 10<sup>-7</sup> |
| Pre-FEV<sub>1</sub>/FVC | 10 | 10:69631004 | 69631004 | A | G | 0.028 | -5.4903 | 1.10 | 6.68 x 10<sup>-7</sup> |
| Pre-FEV<sub>1</sub>/FVC | 10 | 10:69655216 | 69655216 | T | C | 0.052 | -3.9421 | 0.791 | 6.20 x 10<sup>-7</sup> |
| Pre-FEV<sub>1</sub>/FVC | 10 | 10:69655466 | 69655466 | A | G | 0.028 | -5.4903 | 1.10 | 6.68 x 10<sup>-7</sup> |
| Pre-FEV<sub>1</sub>/FVC | 10 | 10:69668476 | 69668476 | A | T | 0.029 | -5.1320 | 1.07 | 1.52 x 10<sup>-6</sup> |
| Pre-FEV<sub>1</sub>/FVC | 10 | 10:69668477 | 69668477 | T | G | 0.028 | -5.4554 | 1.11 | 9.05 x 10<sup>-7</sup> |
| Pre-FEV<sub>1</sub>/FVC | 10 | 10:69673174 | 69673174 | T | C | 0.028 | -5.5138 | 1.14 | 1.21 x 10<sup>-6</sup> |
| Pre-FEV<sub>1</sub>/FVC | 10 | 10:69693013 | 69693013 | C | A | 0.051 | -3.9993 | 0.797 | 5.25 x 10<sup>-7</sup> |
| Pre-FEV<sub>1</sub>/FVC | 10 | 10:69699710 | 69699710 | A | G | 0.051 | -4.1297 | 0.800 | 2.41 x 10<sup>-7</sup> |
| Pre-FEV<sub>1</sub>/FVC | 10 | 10:69701403 | 69701403 | T | C | 0.051 | -4.1297 | 0.800 | 2.41 x 10<sup>-7</sup> |
| Pre-FEV<sub>1</sub>/FVC | 10 | 10:69702559 | 69702559 | T | C | 0.051 | -4.1303 | 0.800 | 2.40 x 10<sup>-7</sup> |
| Pre-FEV<sub>1</sub>/FVC | 11 | 11:23684142 | 23684142 | G | C | 0.011 | -8.5207 | 1.76 | 1.30 x 10<sup>-6</sup> |
| Pre-FEV<sub>1</sub>/FVC | 11 | 11:116627541 | 116627541 | T | C | 0.013 | -8.3846 | 1.67 | 5.49 x 10<sup>-7</sup> |
| Pre-FEV<sub>1</sub>/FVC | 13 | 13:26171336 | 26171336 | C | T | 0.012 | -8.8936 | 1.82 | 1.09 x 10<sup>-6</sup> |
| Pre-FEV<sub>1</sub>/FVC | 13 | 13:26172491 | 26172491 | G | A | 0.012 | -8.8941 | 1.82 | 1.09 x 10<sup>-6</sup> |
| Pre-FEV<sub>1</sub>/FVC | 13 | 13:26173637 | 26173637 | C | G | 0.012 | -8.8936 | 1.82 | 1.09 x 10<sup>-6</sup> |
| Pre-FEV<sub>1</sub>/FVC | 13 | 13:26178584 | 26178584 | G | T | 0.012 | -9.2967 | 1.82 | 3.35 x 10<sup>-7</sup> |
| Pre-FEV<sub>1</sub>/FVC | 13 | 13:26179307 | 26179307 | A | G | 0.012 | -9.4254 | 1.86 | 3.97 x 10<sup>-7</sup> |
| Pre-FEV<sub>1</sub>/FVC | 13 | 13:26179851 | 26179851 | G | A | 0.012 | -9.4264 | 1.86 | 3.97 x 10<sup>-7</sup> |
| Pre-FEV<sub>1</sub>/FVC | 13 | 13:26180215 | 26180215 | C | G | 0.012 | -9.4264 | 1.86 | 3.97 x 10<sup>-7</sup> |
| Pre-FEV<sub>1</sub>/FVC | 13 | 13:26180560 | 26180560 | T | C | 0.012 | -9.4272 | 1.86 | 3.96 x 10<sup>-7</sup> |
| Pre-FEV<sub>1</sub>/FVC | 13 | 13:26180708 | 26180708 | C | A | 0.012 | -9.4264 | 1.86 | 3.97 x 10<sup>-7</sup> |
| Pre-FEV<sub>1</sub>/FVC | 13 | 13:26180788 | 26180788 | C | T | 0.012 | -9.4264 | 1.86 | 3.97 x 10<sup>-7</sup> |
| Pre-FEV<sub>1</sub>/FVC | 13 | 13:26181052 | 26181052 | A | T | 0.012 | -9.4264 | 1.86 | 3.97 x 10<sup>-7</sup> |
| Pre-FEV<sub>1</sub>/FVC | 13 | 13:26181377 | 26181377 | T | A | 0.012 | -9.4264 | 1.86 | 3.97 x 10<sup>-7</sup> |
| Pre-FEV<sub>1</sub>/FVC | 13 | 13:26181548 | 26181548 | T | C | 0.012 | -9.4264 | 1.86 | 3.97 x 10<sup>-7</sup> |
| Pre-FEV<sub>1</sub>/FVC | 13 | 13:26181808 | 26181808 | T | C | 0.012 | -9.4264 | 1.86 | 3.97 x 10<sup>-7</sup> |
| Pre-FEV<sub>1</sub>/FVC | 13 | 13:26182111 | 26182111 | G | A | 0.012 | -9.4264 | 1.86 | 3.97 x 10<sup>-7</sup> |
| Pre-FEV<sub>1</sub>/FVC | 13 | 13:26186904 | 26186904 | C | T | 0.012 | -9.4272 | 1.86 | 3.96 x 10<sup>-7</sup> |
| Pre-FEV<sub>1</sub>/FVC | 13 | 13:26187050 | 26187050 | T | C | 0.012 | -9.4264 | 1.86 | 3.97 x 10<sup>-7</sup> |
| Pre-FEV<sub>1</sub>/FVC | 13 | 13:26187226 | 26187226 | G | A | 0.012 | -9.4272 | 1.86 | 3.96 x 10<sup>-7</sup> |
| Pre-FEV<sub>1</sub>/FVC | 13 | 13:26187654 | 26187654 | G | A | 0.012 | -9.4264 | 1.86 | 3.97 x 10<sup>-7</sup> |
| Pre-FEV<sub>1</sub>/FVC | 13 | 13:26190577 | 26190577 | T | C | 0.011 | -10.0394 | 1.90 | 1.23 x 10<sup>-7</sup> |
| Pre-FEV<sub>1</sub>/FVC | 13 | 13:26191329 | 26191329 | T | C | 0.012 | -9.4272 | 1.86 | 3.96 x 10<sup>-7</sup> |
| Pre-FEV<sub>1</sub>/FVC | 13 | 13:26191597 | 26191597 | G | T | 0.012 | -9.4272 | 1.86 | 3.96 x 10<sup>-7</sup> |
| Pre-FEV<sub>1</sub>/FVC | 13 | 13:26198053 | 26198053 | T | G | 0.011 | -9.7840 | 1.90 | 2.52 x 10<sup>-7</sup> |
| Pre-FEV<sub>1</sub>/FVC | 13 | 13:26201829 | 26201829 | G | C | 0.011 | -9.7840 | 1.90 | 2.52 x 10<sup>-7</sup> |
| Pre-FEV<sub>1</sub>/FVC | 13 | 13:26235394 | 26235394 | G | A | 0.010 | -10.5334 | 1.94 | 5.80 x 10<sup>-8</sup> |
| Pre-FEV<sub>1</sub>/FVC | 13 | 13:26247080 | 26247080 | G | A | 0.010 | -10.5334 | 1.94 | 5.80 x 10<sup>-8</sup> |
| Pre-FEV<sub>1</sub>/FVC | 13 | 13:26262378 | 26262378 | T | G | 0.011 | -10.5325 | 1.94 | 5.81 x 10<sup>-8</sup> |
| Pre-FEV<sub>1</sub>/FVC | 13 | 13:26268604 | 26268604 | A | C | 0.011 | -10.5325 | 1.94 | 5.81 x 10<sup>-8</sup> |
| Pre-FEV<sub>1</sub>/FVC | 13 | 13:26280122 | 26280122 | A | G | 0.011 | -9.5140 | 1.90 | 5.52 x 10<sup>-7</sup> |
| Pre-FEV<sub>1</sub>/FVC | 14 | 14:94463882 | 94463882 | C | T | 0.028 | -5.9224 | 1.23 | 1.35 x 10<sup>-6</sup> |
| Pre-FEV<sub>1</sub>/FVC | 14 | 14:94467860 | 94467860 | T | C | 0.028 | -5.9224 | 1.23 | 1.35 x 10<sup>-6</sup> |
| Pre-FEV<sub>1</sub>/FVC | 14 | 14:94472407 | 94472407 | G | A | 0.036 | -5.5247 | 1.13 | 9.25 x 10<sup>-7</sup> |
| Pre-FEV<sub>1</sub>/FVC | 14 | 14:94472871 | 94472871 | G | T | 0.036 | -5.4523 | 1.12 | 1.10 x 10<sup>-6</sup> |
| Pre-FEV<sub>1</sub>/FVC | 14 | 14:94479922 | 94479922 | A | G | 0.033 | -5.8213 | 1.15 | 3.74 x 10<sup>-7</sup> |
| Pre-FEV<sub>1</sub>/FVC | 15 | 15:78801396 | 78801396 | C | A | 0.139 | -2.6943 | 0.560 | 1.53 x 10<sup>-6</sup> |
| Pre-FEV<sub>1</sub>/FVC | 15 | 15:78805789 | 78805789 | C | G | 0.140 | -2.7840 | 0.559 | 6.37 x 10<sup>-7</sup> |
| Pre-FEV<sub>1</sub>/FVC | 15 | 15:78807732 | 78807732 | A | C | 0.140 | -2.7840 | 0.559 | 6.37 x 10<sup>-7</sup> |
| Pre-FEV<sub>1</sub>/FVC | 15 | 15:78811677 | 78811677 | C | T | 0.140 | -2.7233 | 0.560 | 1.14 x 10<sup>-6</sup> |
| Pre-FEV<sub>1</sub>/FVC | 15 | 15:78821368 | 78821368 | A | C | 0.139 | -2.6730 | 0.561 | 1.90 x 10<sup>-6</sup> |
| Pre-FEV<sub>1</sub>/FVC | 15 | 15:93817218 | 93817218 | A | G | 0.034 | -5.0102 | 1.01 | 7.11 x 10<sup>-7</sup> |
| Pre-FEV<sub>1</sub>/FVC | 15 | 15:93818088 | 93818088 | G | C | 0.035 | -5.0107 | 1.01 | 7.10 x 10<sup>-7</sup> |
| Pre-FEV<sub>1</sub>/FVC | 16 | 16:19508306 | 19508306 | A | G | 0.062 | -3.8078 | 0.786 | 1.25 x 10<sup>-6</sup> |
| Pre-FEV<sub>1</sub>/FVC | 16 | 16:19597633 | 19597633 | C | G | 0.043 | -4.5022 | 0.945 | 1.88 x 10<sup>-6</sup> |
| Pre-FEV<sub>1</sub>/FVC | 18 | 18:55422267 | 55422267 | A | G | 0.013 | -7.9629 | 1.67 | 1.80 x 10<sup>-6</sup> |

**Supplementary Table 6:**
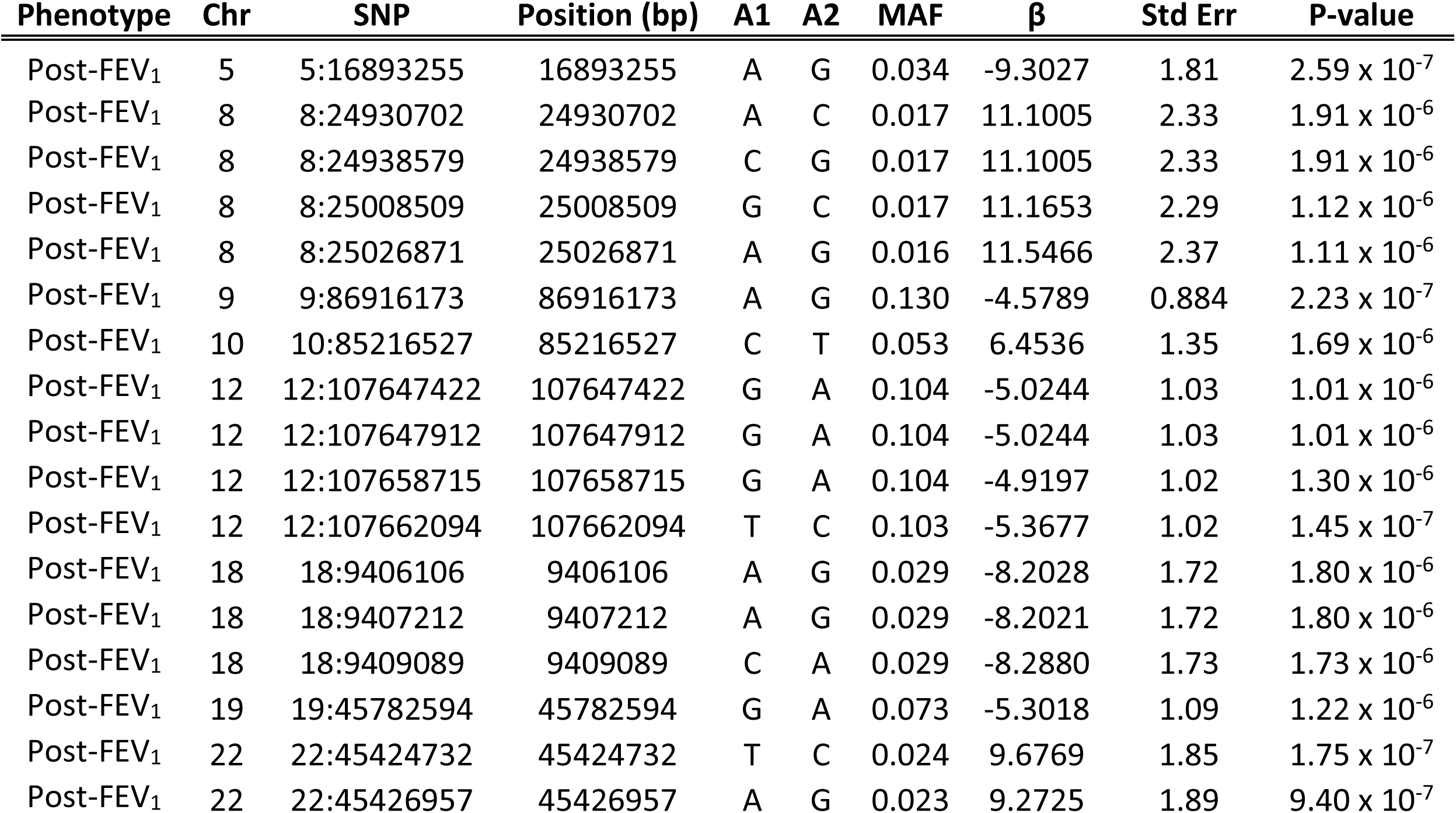
Suggestive associations with Post-FEV_1_.

**Supplementary Table 7:** Suggestive associations with Post-FVC.

| Phenotype | Chr | SNP | Position (bp) | A1 | A2 | MAF | $\beta$ | Std Err | P-value |
| --- | --- | --- | --- | --- | --- | --- | --- | --- | --- |
| Post-FVC | 1 | 1:195763875 | 195763875 | C | T | 0.255 | -3.3983 | 0.700 | $1.21 \times 10^{-6}$ |
| Post-FVC | 1 | 1:195771663 | 195771663 | T | C | 0.259 | -3.4962 | 0.698 | $5.52 \times 10^{-7}$ |
| Post-FVC | 1 | 1:195771841 | 195771841 | G | A | 0.259 | -3.4962 | 0.698 | $5.52 \times 10^{-7}$ |
| Post-FVC | 1 | 1:195778940 | 195778940 | A | T | 0.246 | -3.6116 | 0.720 | $5.29 \times 10^{-7}$ |
| Post-FVC | 2 | 2:115966363 | 115966363 | C | T | 0.072 | 6.0241 | 1.26 | $1.61 \times 10^{-6}$ |
| Post-FVC | 2 | 2:115978242 | 115978242 | C | T | 0.071 | 6.1979 | 1.27 | $9.64 \times 10^{-7}$ |
| Post-FVC | 2 | 2:115982589 | 115982589 | C | T | 0.074 | 6.0001 | 1.24 | $1.37 \times 10^{-6}$ |
| Post-FVC | 6 | 6:162713288 | 162713288 | A | G | 0.014 | 14.0917 | 2.68 | $1.45 \times 10^{-7}$ |
| Post-FVC | 8 | 8:83741379 | 83741379 | A | C | 0.094 | -5.0755 | 1.07 | $1.89 \times 10^{-6}$ |
| Post-FVC | 8 | 8:119895298 | 119895298 | C | A | 0.021 | 11.6893 | 2.31 | $4.31 \times 10^{-7}$ |
| Post-FVC | 11 | 11:118902275 | 118902275 | C | T | 0.010 | 17.9775 | 3.29 | $4.83 \times 10^{-8}$ |
| Post-FVC | 11 | 11:118905095 | 118905095 | A | G | 0.010 | 17.9775 | 3.29 | $4.83 \times 10^{-8}$ |
| Post-FVC | 11 | 11:118905316 | 118905316 | T | C | 0.010 | 17.9775 | 3.29 | $4.83 \times 10^{-8}$ |
| Post-FVC | 11 | 11:118906065 | 118906065 | A | G | 0.010 | 17.9775 | 3.29 | $4.83 \times 10^{-8}$ |
| Post-FVC | 11 | 11:118906240 | 118906240 | T | C | 0.010 | 17.9775 | 3.29 | $4.83 \times 10^{-8}$ |
| Post-FVC | 11 | 11:118906745 | 118906745 | C | G | 0.010 | 17.9775 | 3.29 | $4.83 \times 10^{-8}$ |
| Post-FVC | 11 | 11:118907923 | 118907923 | G | T | 0.010 | 18.2978 | 3.41 | $7.85 \times 10^{-8}$ |
| Post-FVC | 11 | 11:118932913 | 118932913 | C | T | 0.010 | 18.9094 | 3.29 | $9.47 \times 10^{-9}$ |
| Post-FVC | 12 | 12:107662094 | 107662094 | T | C | 0.103 | -5.1924 | 1.05 | $6.91 \times 10^{-7}$ |
| Post-FVC | 12 | 12:115797499 | 115797499 | G | A | 0.060 | 6.6259 | 1.33 | $6.34 \times 10^{-7}$ |
| Post-FVC | 17 | 17:74377172 | 74377172 | T | C | 0.110 | 4.8181 | 0.955 | $4.54 \times 10^{-7}$ |
| Post-FVC | 18 | 18:67404762 | 67404762 | A | C | 0.010 | 16.4840 | 3.40 | $1.28 \times 10^{-6}$ |
| Post-FVC | 18 | 18:67404973 | 67404973 | A | G | 0.010 | 16.4840 | 3.40 | $1.28 \times 10^{-6}$ |
| Post-FVC | 18 | 18:67404990 | 67404990 | T | G | 0.010 | 16.4840 | 3.40 | $1.28 \times 10^{-6}$ |
| Post-FVC | 19 | 19:4289259 | 4289259 | T | C | 0.106 | 5.1672 | 0.967 | $9.21 \times 10^{-8}$ |
| Post-FVC | 19 | 19:4291817 | 4291817 | T | C | 0.111 | 5.2674 | 0.963 | $4.52 \times 10^{-8}$ |
| Post-FVC | 19 | 19:4294011 | 4294011 | C | A | 0.140 | 4.3027 | 0.885 | $1.15 \times 10^{-6}$ |
| Post-FVC | 19 | 19:4294410 | 4294410 | C | T | 0.157 | 4.0184 | 0.829 | $1.25 \times 10^{-6}$ |
| Post-FVC | 19 | 19:4298416 | 4298416 | A | G | 0.153 | 4.1478 | 0.847 | $9.85 \times 10^{-7}$ |
| Post-FVC | 19 | 19:4302569 | 4302569 | G | A | 0.156 | 4.0985 | 0.840 | $1.05 \times 10^{-6}$ |
| Post-FVC | 19 | 19:36372775 | 36372775 | T | C | 0.129 | -4.2774 | 0.889 | $1.49 \times 10^{-6}$ |
| Post-FVC | 19 | 19:36374386 | 36374386 | G | T | 0.272 | -3.3295 | 0.698 | $1.87 \times 10^{-6}$ |
| Post-FVC | 19 | 19:36376980 | 36376980 | A | G | 0.129 | -4.2471 | 0.890 | $1.82 \times 10^{-6}$ |
| Post-FVC | 21 | 21:18234328 | 18234328 | T | C | 0.012 | 12.7085 | 2.62 | $1.19 \times 10^{-6}$ |
| Post-FVC | 21 | 21:18234404 | 18234404 | A | G | 0.012 | 12.7085 | 2.62 | $1.19 \times 10^{-6}$ |
| Post-FVC | 21 | 21:43755177 | 43755177 | G | A | 0.091 | 5.5767 | 1.15 | $1.25 \times 10^{-6}$ |

**Supplementary Table 8:** Suggestive associations with Post-FEV_1_/FVC.

| Phenotype | Chr | SNP | Position (bp) | A1 | A2 | MAF | $\beta$ | Std Err | P-value |
| --- | --- | --- | --- | --- | --- | --- | --- | --- | --- |
| Post-FEV<sub>1</sub>/FVC | 1 | 1:117580100 | 117580100 | G | A | 0.015 | -7.0892 | 1.45 | 1.01 x 10<sup>-6</sup> |
| Post-FEV<sub>1</sub>/FVC | 1 | 1:117580363 | 117580363 | C | T | 0.015 | -7.0892 | 1.45 | 1.01 x 10<sup>-6</sup> |
| Post-FEV<sub>1</sub>/FVC | 1 | 1:180886944 | 180886944 | C | T | 0.015 | -7.0382 | 1.35 | 1.76 x 10<sup>-7</sup> |
| Post-FEV<sub>1</sub>/FVC | 1 | 1:180959189 | 180959189 | G | A | 0.010 | -7.7534 | 1.58 | 9.13 x 10<sup>-7</sup> |
| Post-FEV<sub>1</sub>/FVC | 1 | 1:180970045 | 180970045 | C | T | 0.010 | -7.7534 | 1.58 | 9.13 x 10<sup>-7</sup> |
| Post-FEV<sub>1</sub>/FVC | 1 | 1:185527158 | 185527158 | G | C | 0.013 | -9.4499 | 1.86 | 3.80 x 10<sup>-7</sup> |
| Post-FEV<sub>1</sub>/FVC | 2 | 2:26387193 | 26387193 | G | C | 0.125 | -2.4299 | 0.510 | 1.90 x 10<sup>-6</sup> |
| Post-FEV<sub>1</sub>/FVC | 2 | 2:26390965 | 26390965 | A | G | 0.131 | -2.4049 | 0.504 | 1.79 x 10<sup>-6</sup> |
| Post-FEV<sub>1</sub>/FVC | 2 | 2:26392354 | 26392354 | A | C | 0.131 | -2.4220 | 0.504 | 1.56 x 10<sup>-6</sup> |
| Post-FEV<sub>1</sub>/FVC | 2 | 2:171008016 | 171008016 | G | A | 0.014 | -6.9043 | 1.37 | 4.70 x 10<sup>-7</sup> |
| Post-FEV<sub>1</sub>/FVC | 3 | 3:22321021 | 22321021 | A | G | 0.013 | -7.8568 | 1.52 | 2.17 x 10<sup>-7</sup> |
| Post-FEV<sub>1</sub>/FVC | 3 | 3:49908748 | 49908748 | A | G | 0.012 | -8.5469 | 1.70 | 5.07 x 10<sup>-7</sup> |
| Post-FEV<sub>1</sub>/FVC | 3 | 3:49908976 | 49908976 | C | T | 0.012 | -8.5469 | 1.70 | 5.07 x 10<sup>-7</sup> |
| Post-FEV<sub>1</sub>/FVC | 3 | 3:49945028 | 49945028 | G | A | 0.010 | -9.0560 | 1.86 | 1.11 x 10<sup>-6</sup> |
| Post-FEV<sub>1</sub>/FVC | 4 | 4:55555416 | 55555416 | G | A | 0.031 | -4.4167 | 0.923 | 1.69 x 10<sup>-6</sup> |
| Post-FEV<sub>1</sub>/FVC | 4 | 4:55568875 | 55568875 | A | G | 0.030 | -4.4141 | 0.922 | 1.70 x 10<sup>-6</sup> |
| Post-FEV<sub>1</sub>/FVC | 4 | 4:70284767 | 70284767 | A | T | 0.010 | -9.5784 | 1.87 | 2.89 x 10<sup>-7</sup> |
| Post-FEV<sub>1</sub>/FVC | 4 | 4:70338226 | 70338226 | A | G | 0.012 | -8.7913 | 1.71 | 2.53 x 10<sup>-7</sup> |
| Post-FEV<sub>1</sub>/FVC | 5 | 5:89139642 | 89139642 | G | A | 0.011 | -8.3271 | 1.65 | 4.85 x 10<sup>-7</sup> |
| Post-FEV<sub>1</sub>/FVC | 5 | 5:89174965 | 89174965 | T | C | 0.011 | -8.3271 | 1.65 | 4.85 x 10<sup>-7</sup> |
| Post-FEV<sub>1</sub>/FVC | 5 | 5:89201706 | 89201706 | T | C | 0.011 | -8.3265 | 1.65 | 4.86 x 10<sup>-7</sup> |
| Post-FEV<sub>1</sub>/FVC | 8 | 8:4110597 | 4110597 | A | G | 0.066 | -3.3772 | 0.704 | 1.60 x 10<sup>-6</sup> |
| Post-FEV<sub>1</sub>/FVC | 10 | 10:69583770 | 69583770 | T | C | 0.030 | -4.7114 | 0.982 | 1.59 x 10<sup>-6</sup> |
| Post-FEV<sub>1</sub>/FVC | 11 | 11:35570480 | 35570480 | A | G | 0.016 | -6.1461 | 1.27 | 1.20 x 10<sup>-6</sup> |
| Post-FEV<sub>1</sub>/FVC | 11 | 11:35573791 | 35573791 | T | A | 0.017 | -6.1468 | 1.27 | 1.20 x 10<sup>-6</sup> |
| Post-FEV<sub>1</sub>/FVC | 11 | 11:35577589 | 35577589 | A | G | 0.016 | -6.2517 | 1.30 | 1.66 x 10<sup>-6</sup> |
| Post-FEV<sub>1</sub>/FVC | 12 | 12:4773488 | 4773488 | C | A | 0.015 | -7.0233 | 1.35 | 1.91 x 10<sup>-7</sup> |
| Post-FEV<sub>1</sub>/FVC | 12 | 12:4773489 | 4773489 | C | G | 0.015 | -7.0233 | 1.35 | 1.91 x 10<sup>-7</sup> |
| Post-FEV<sub>1</sub>/FVC | 12 | 12:4777393 | 4777393 | A | G | 0.015 | -7.0361 | 1.40 | 4.98 x 10<sup>-7</sup> |
| Post-FEV<sub>1</sub>/FVC | 12 | 12:4784518 | 4784518 | T | C | 0.018 | -6.3324 | 1.27 | 6.11 x 10<sup>-7</sup> |
| Post-FEV<sub>1</sub>/FVC | 12 | 12:4784582 | 4784582 | T | G | 0.018 | -6.3324 | 1.27 | 6.11 x 10<sup>-7</sup> |
| Post-FEV<sub>1</sub>/FVC | 12 | 12:4786418 | 4786418 | A | G | 0.018 | -6.1523 | 1.25 | 8.81 x 10<sup>-7</sup> |
| Post-FEV<sub>1</sub>/FVC | 12 | 12:4787767 | 4787767 | G | A | 0.018 | -6.1514 | 1.25 | 8.83 x 10<sup>-7</sup> |
| Post-FEV<sub>1</sub>/FVC | 12 | 12:4788226 | 4788226 | A | G | 0.018 | -6.1531 | 1.25 | 8.80 x 10<sup>-7</sup> |
| Post-FEV<sub>1</sub>/FVC | 12 | 12:4793670 | 4793670 | A | G | 0.020 | -5.7624 | 1.20 | 1.63 x 10<sup>-6</sup> |
| Post-FEV<sub>1</sub>/FVC | 14 | 14:38963901 | 38963901 | C | T | 0.084 | -3.3320 | 0.673 | 7.50 x 10<sup>-7</sup> |
| Post-FEV<sub>1</sub>/FVC | 15 | 15:76052762 | 76052762 | T | G | 0.344 | -1.8331 | 0.372 | 8.07 x 10<sup>-7</sup> |
| Post-FEV<sub>1</sub>/FVC | 15 | 15:76057486 | 76057486 | A | G | 0.336 | -1.8270 | 0.373 | 9.50 x 10<sup>-7</sup> |
| Post-FEV<sub>1</sub>/FVC | 15 | 15:76057493 | 76057493 | T | C | 0.355 | -1.8888 | 0.368 | 2.96 x 10<sup>-7</sup> |
| Post-FEV<sub>1</sub>/FVC | 15 | 15:85798401 | 85798401 | A | G | 0.018 | -6.7846 | 1.27 | 9.05 x 10<sup>-8</sup> |
| Post-FEV<sub>1</sub>/FVC | 15 | 15:85819723 | 85819723 | C | T | 0.020 | -5.8136 | 1.21 | 1.68 x 10<sup>-6</sup> |
| Post-FEV<sub>1</sub>/FVC | 15 | 15:85885031 | 85885031 | A | C | 0.034 | -4.9213 | 0.955 | 2.55 x 10<sup>-7</sup> |
| Post-FEV<sub>1</sub>/FVC | 15 | 15:88957199 | 88957199 | G | T | 0.089 | -2.8154 | 0.591 | 1.93 x 10<sup>-6</sup> |
| Post-FEV<sub>1</sub>/FVC | 15 | 15:93817218 | 93817218 | A | G | 0.034 | -5.0171 | 0.968 | 2.19 x 10<sup>-7</sup> |
| Post-FEV<sub>1</sub>/FVC | 15 | 15:93818088 | 93818088 | G | C | 0.035 | -5.0166 | 0.968 | 2.20 x 10<sup>-7</sup> |
| Post-FEV<sub>1</sub>/FVC | 17 | 17:57263393 | 57263393 | A | G | 0.346 | -1.8380 | 0.380 | 1.30 x 10<sup>-6</sup> |
| Post-FEV<sub>1</sub>/FVC | 17 | 17:57273681 | 57273681 | C | T | 0.345 | -1.8303 | 0.379 | 1.35 x 10<sup>-6</sup> |
| Post-FEV<sub>1</sub>/FVC | 17 | 17:57274481 | 57274481 | C | T | 0.345 | -1.8303 | 0.379 | 1.35 x 10<sup>-6</sup> |
| Post-FEV<sub>1</sub>/FVC | 17 | 17:57276710 | 57276710 | A | G | 0.345 | -1.8301 | 0.379 | 1.35 x 10<sup>-6</sup> |
| Post-FEV<sub>1</sub>/FVC | 17 | 17:78917207 | 78917207 | T | G | 0.015 | -7.8302 | 1.58 | 7.44 x 10<sup>-7</sup> |

**Supplementary Figure 6:**
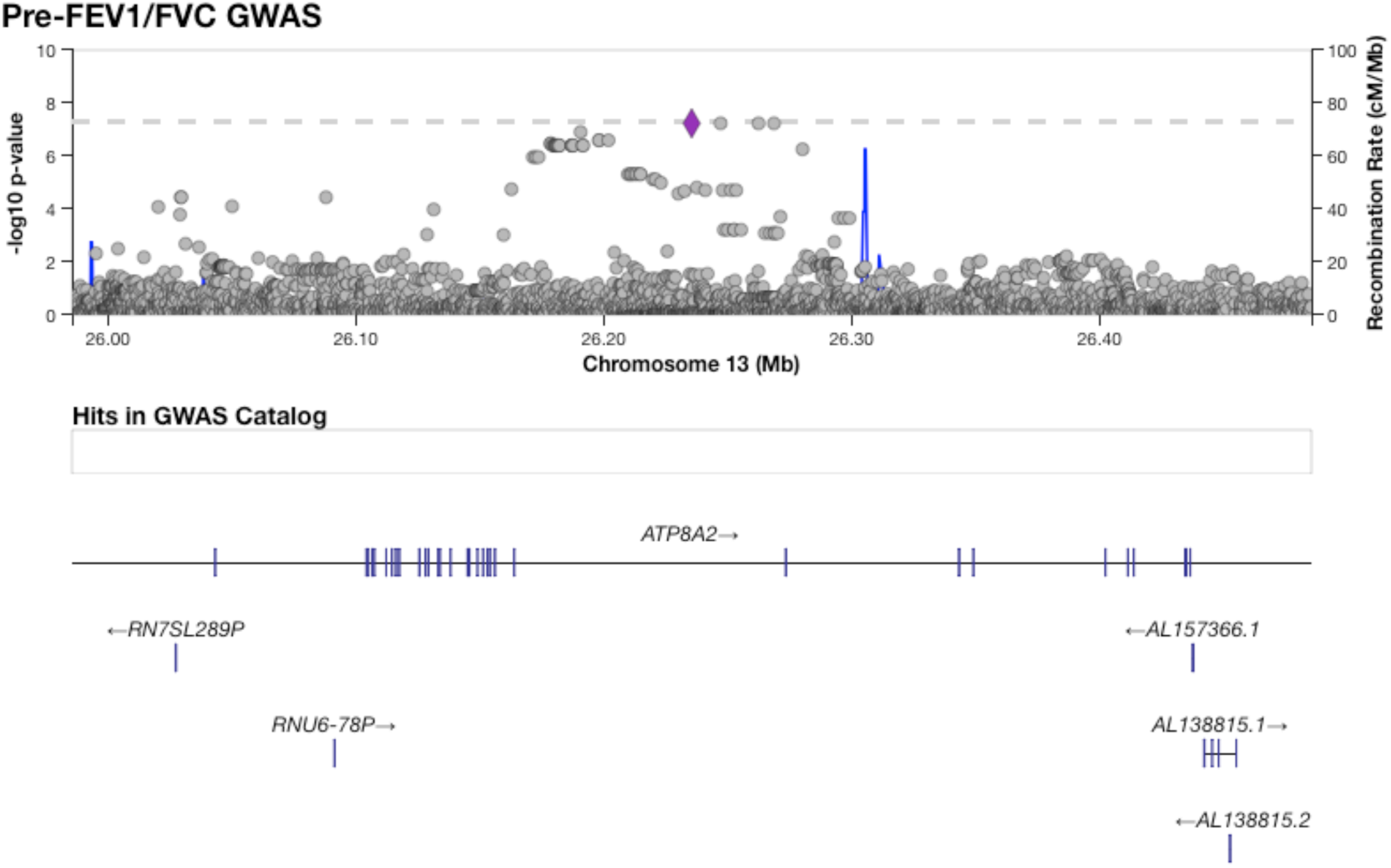
The region encompassing four variants on chromosome 13 significantly associated with Pre-FEV_1_/FVC.

**Supplementary Figure 7:**
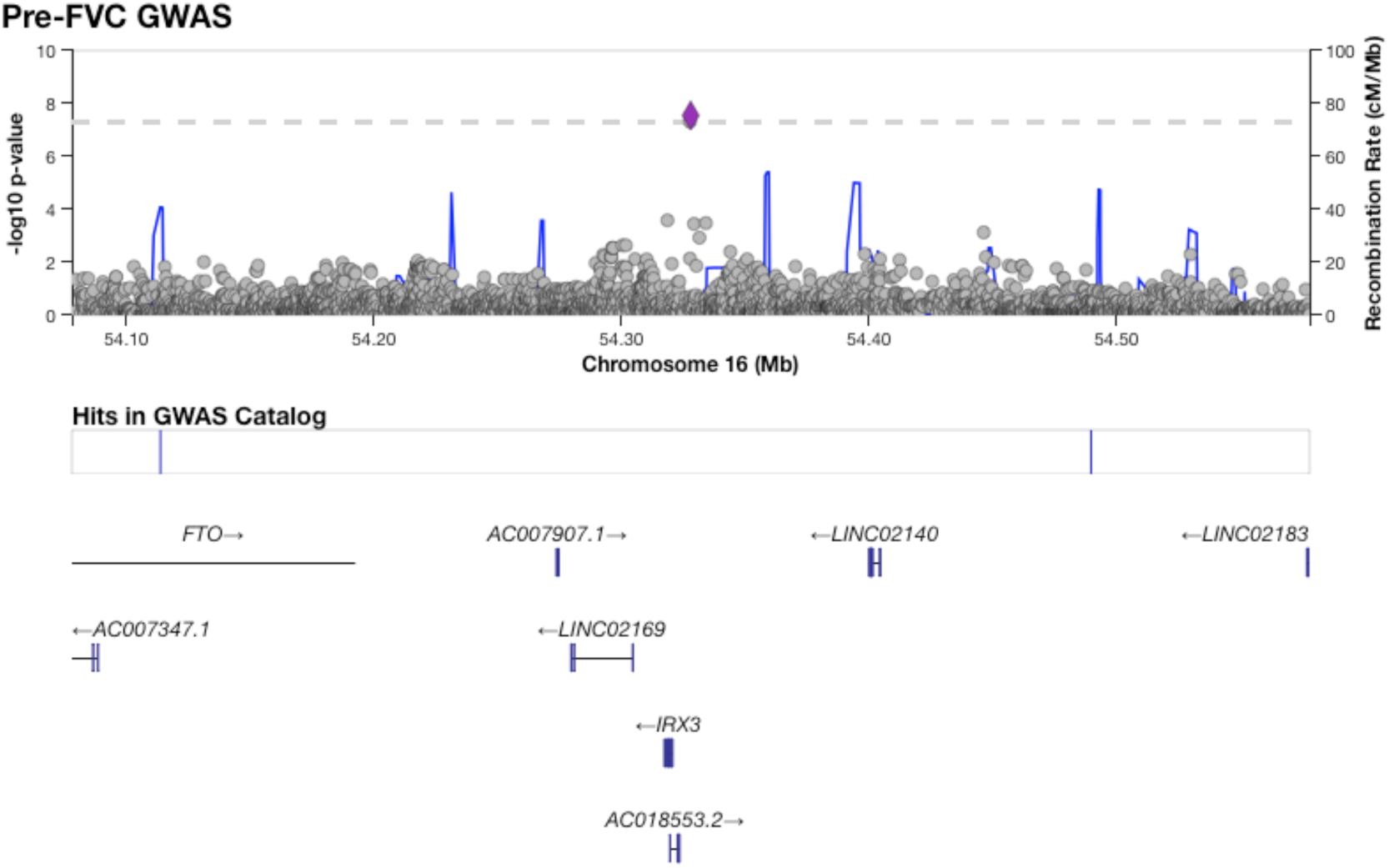
The region encompassing two variants on chromosome 16 significantly associated with Pre-FVC.

**Supplementary Figure 8:**
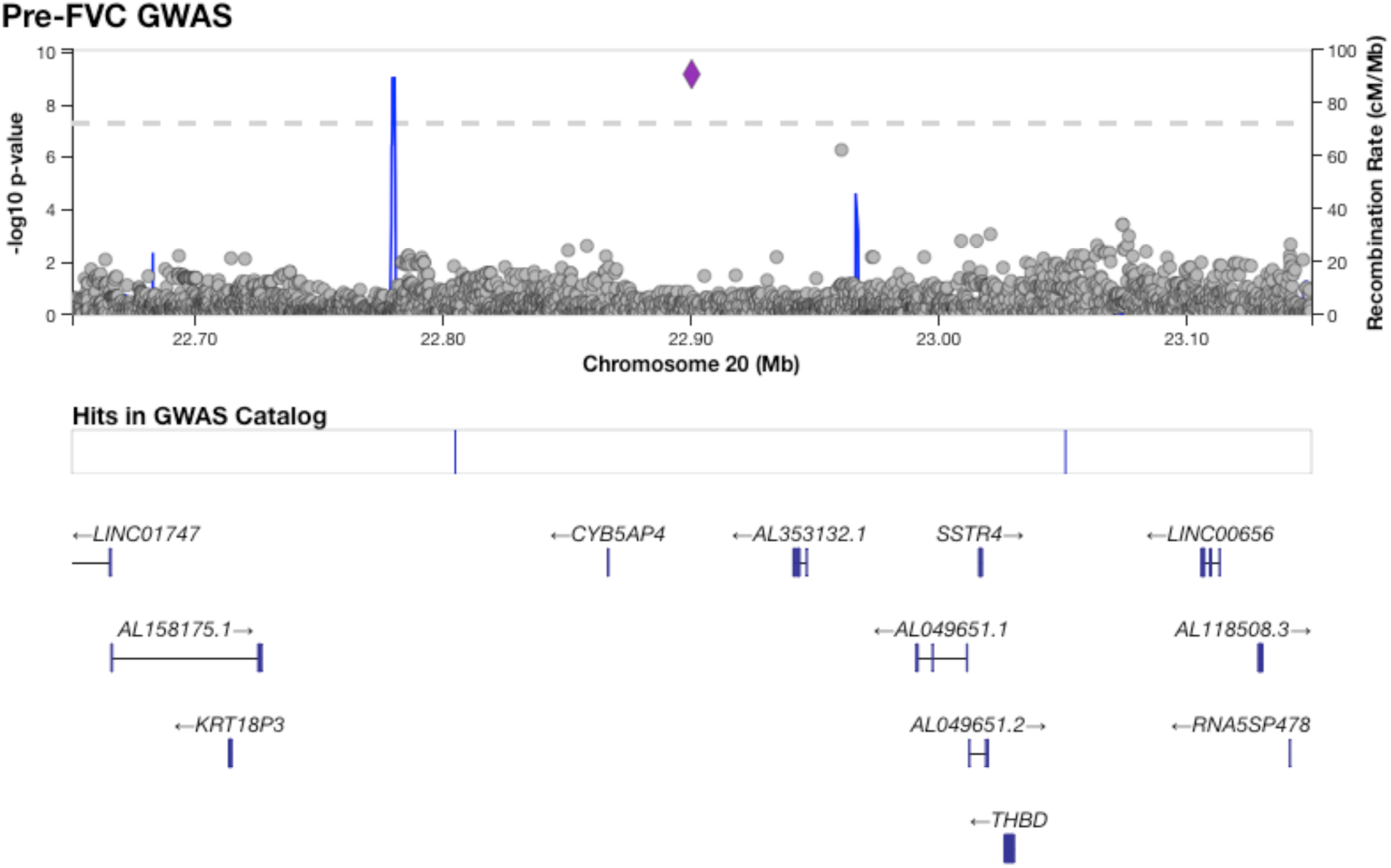
The region encompassing a variant on chromosome 20 significantly associated with Pre-FVC.

**Supplementary Figure 9:**
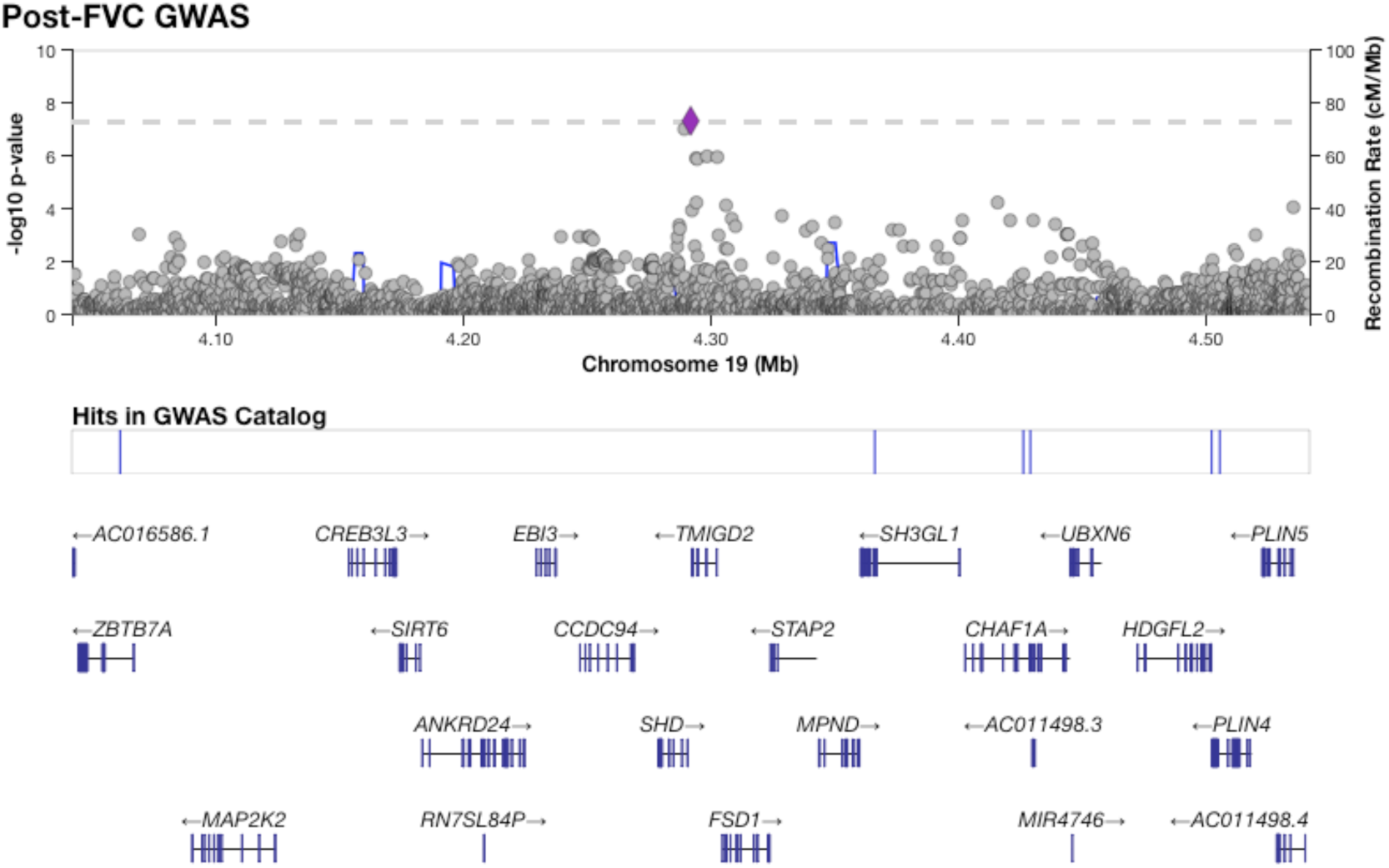
The region encompassing two variants on chromosome 19 significantly associated with Post-FVC.

**Supplementary Figure 10:**
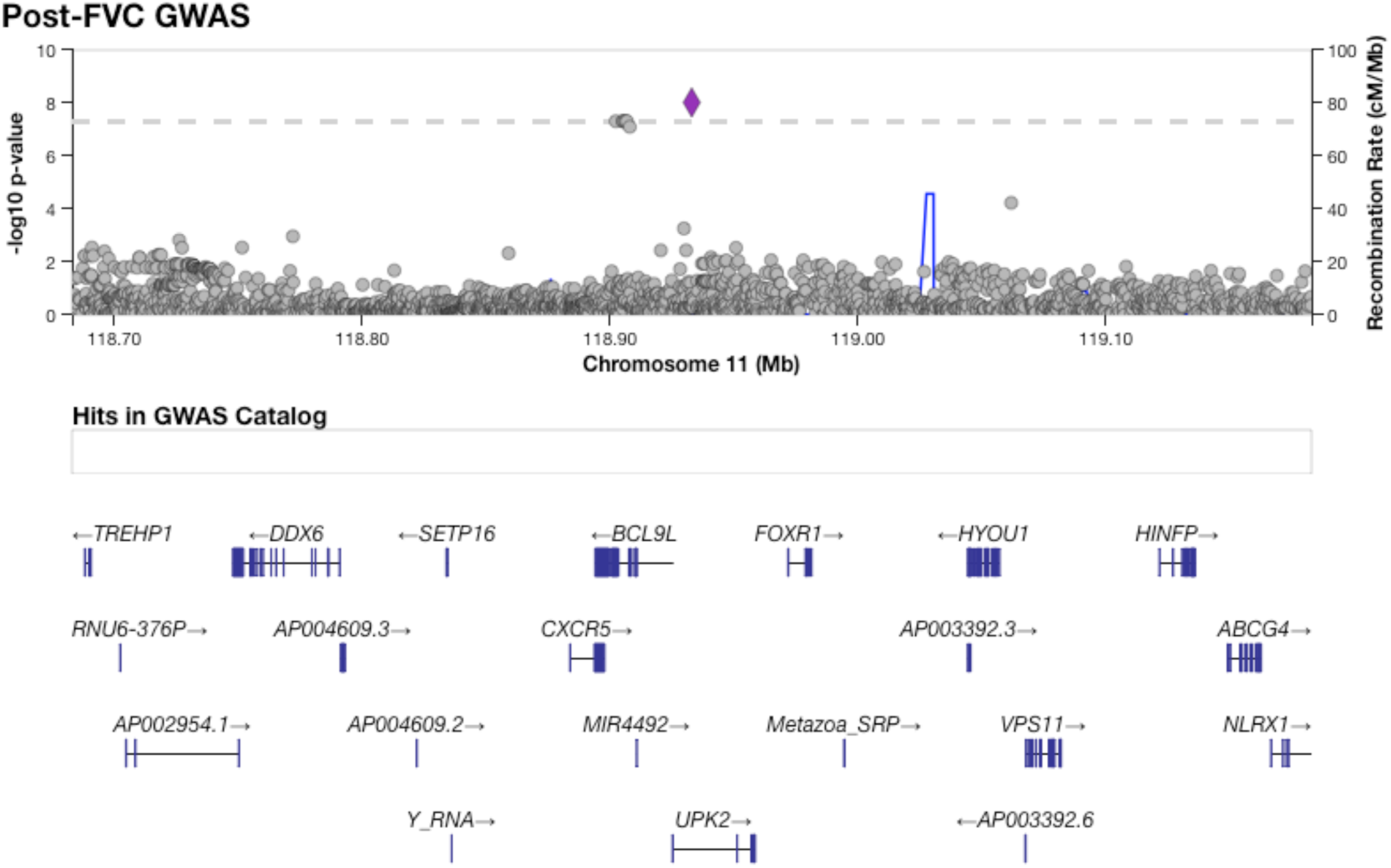
The region encompassing eight variants on chromosome 11 significantly associated with Post-FVC.

**Supplementary Figure 11:**
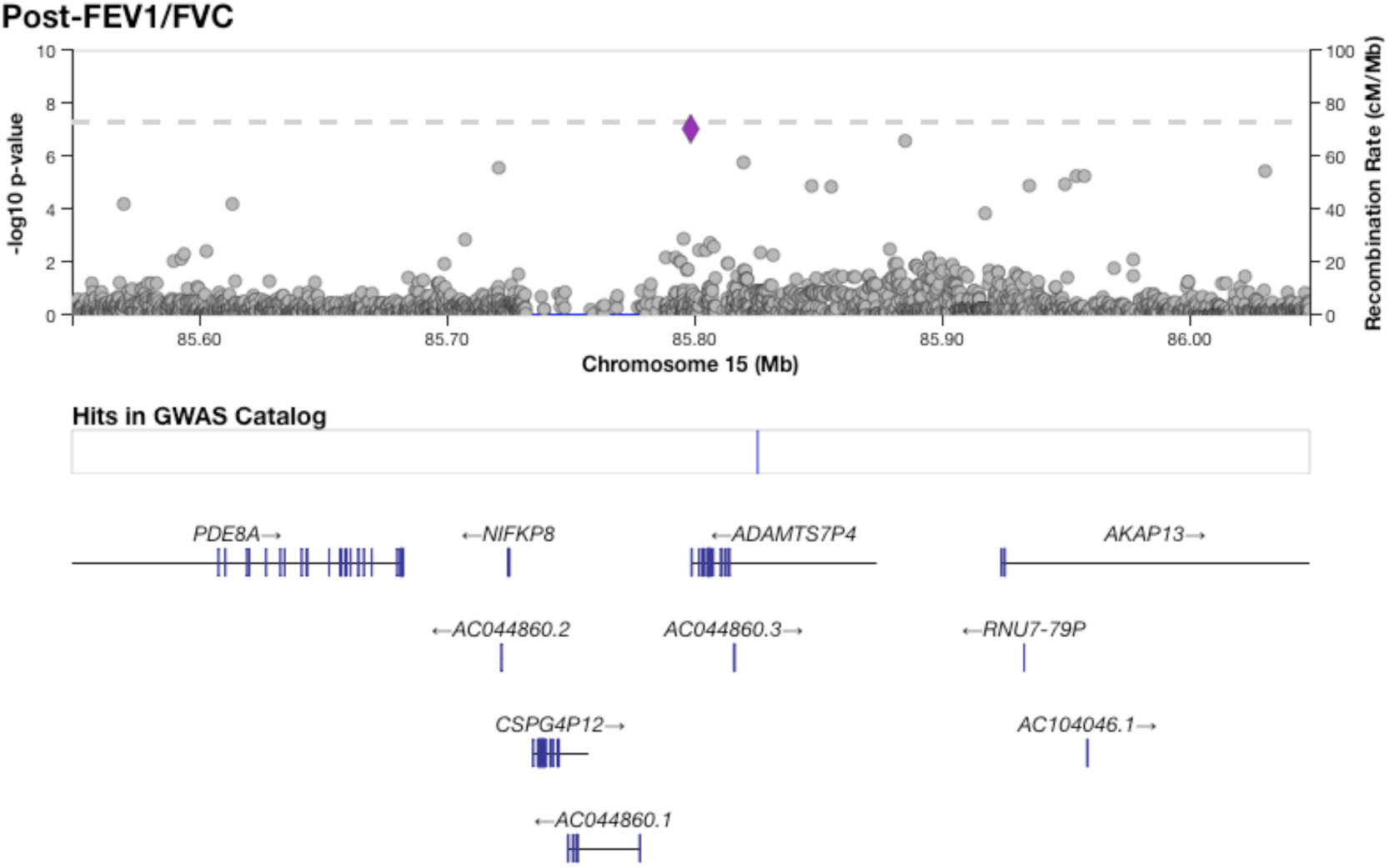
The region surrounding a single variant on chromosome 15 significantly associated with Post-FEV_1_/FVC.

**Supplementary Figure 12:**
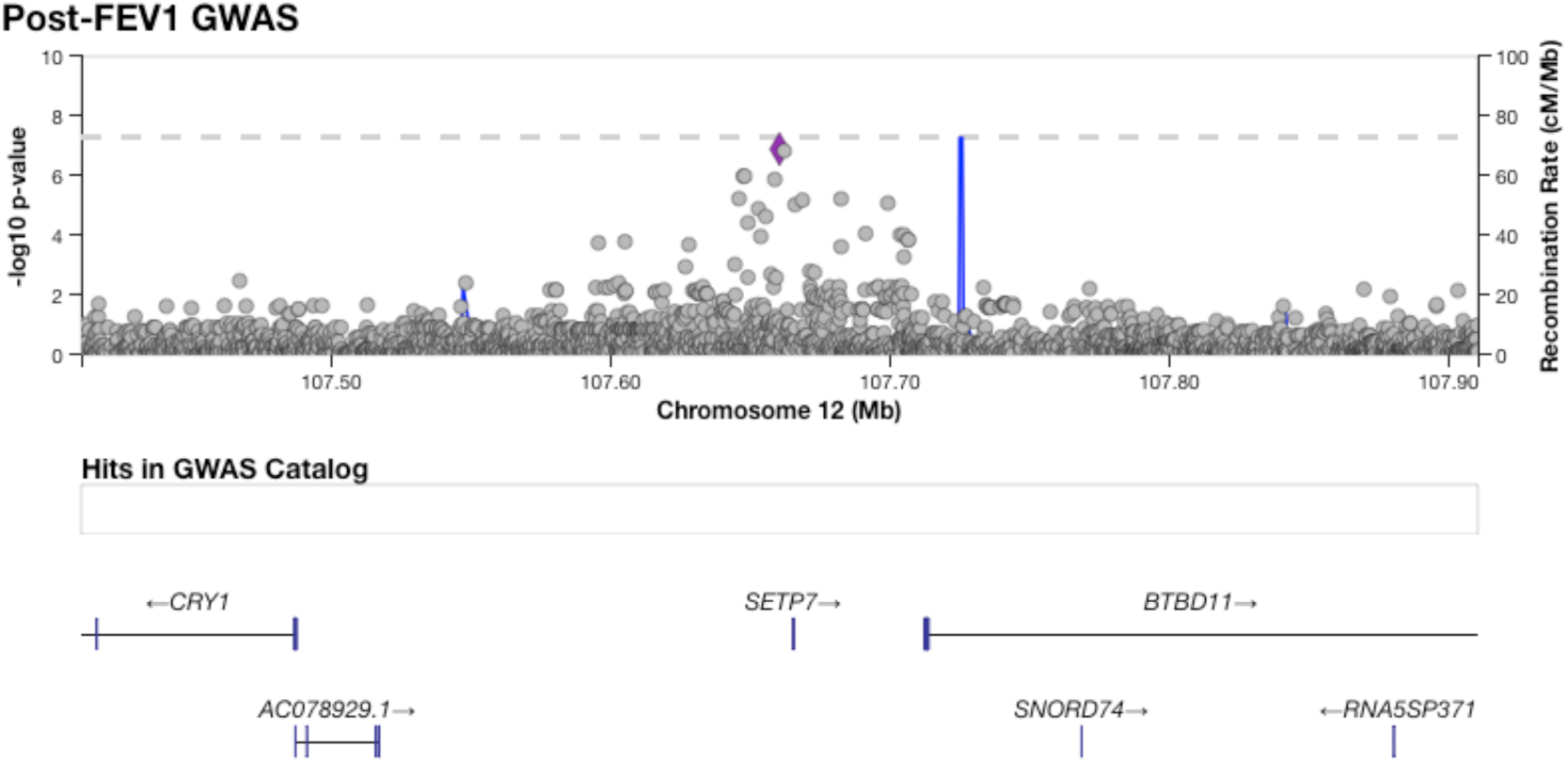
The region around four variants suggestively associated with Post-FEV_1_.

**Supplementary Figure 13:**
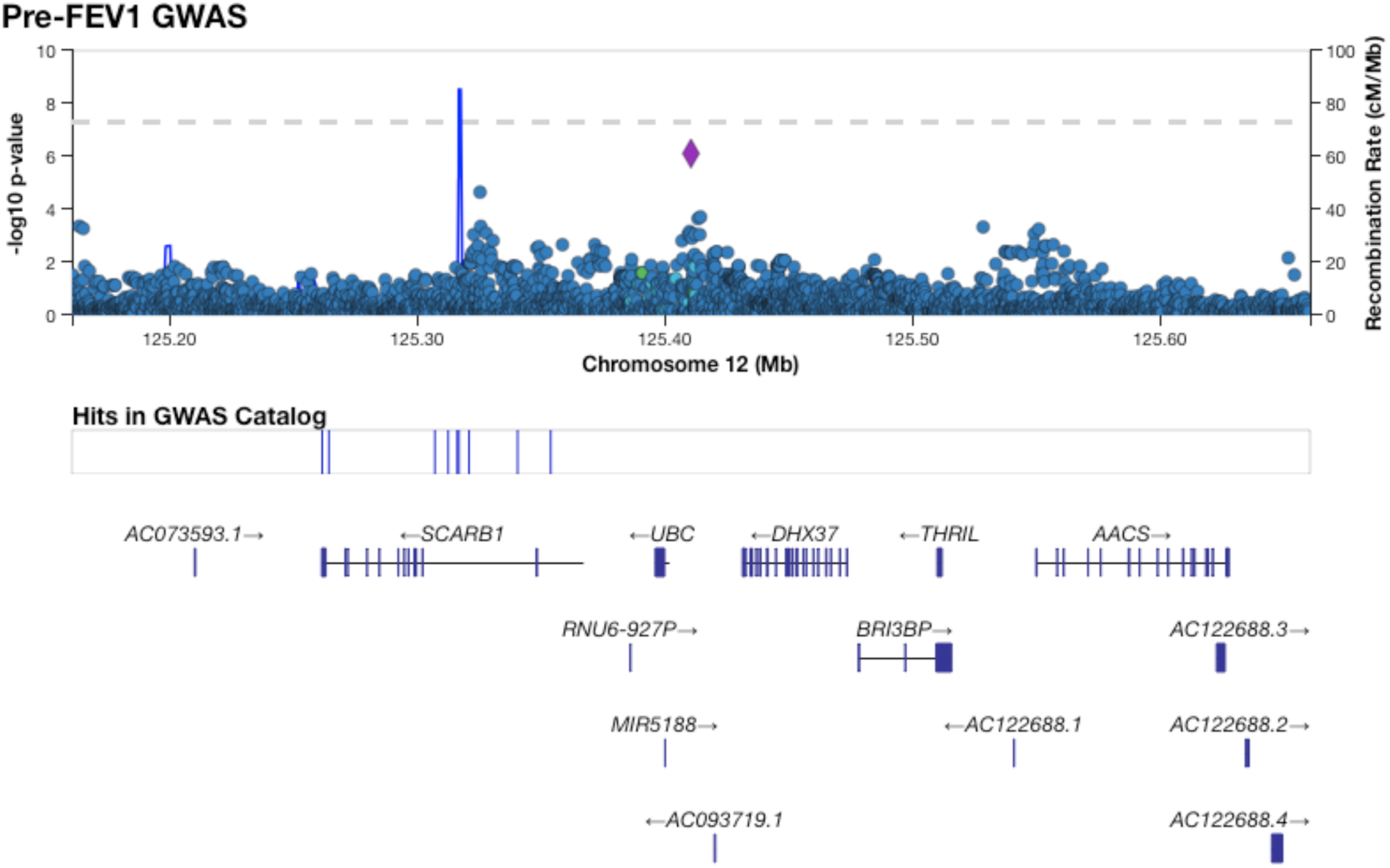
The region around a variant on chromosome 12 suggestively associated with Pre-FEV_1_.

**Supplementary Figure 14:**
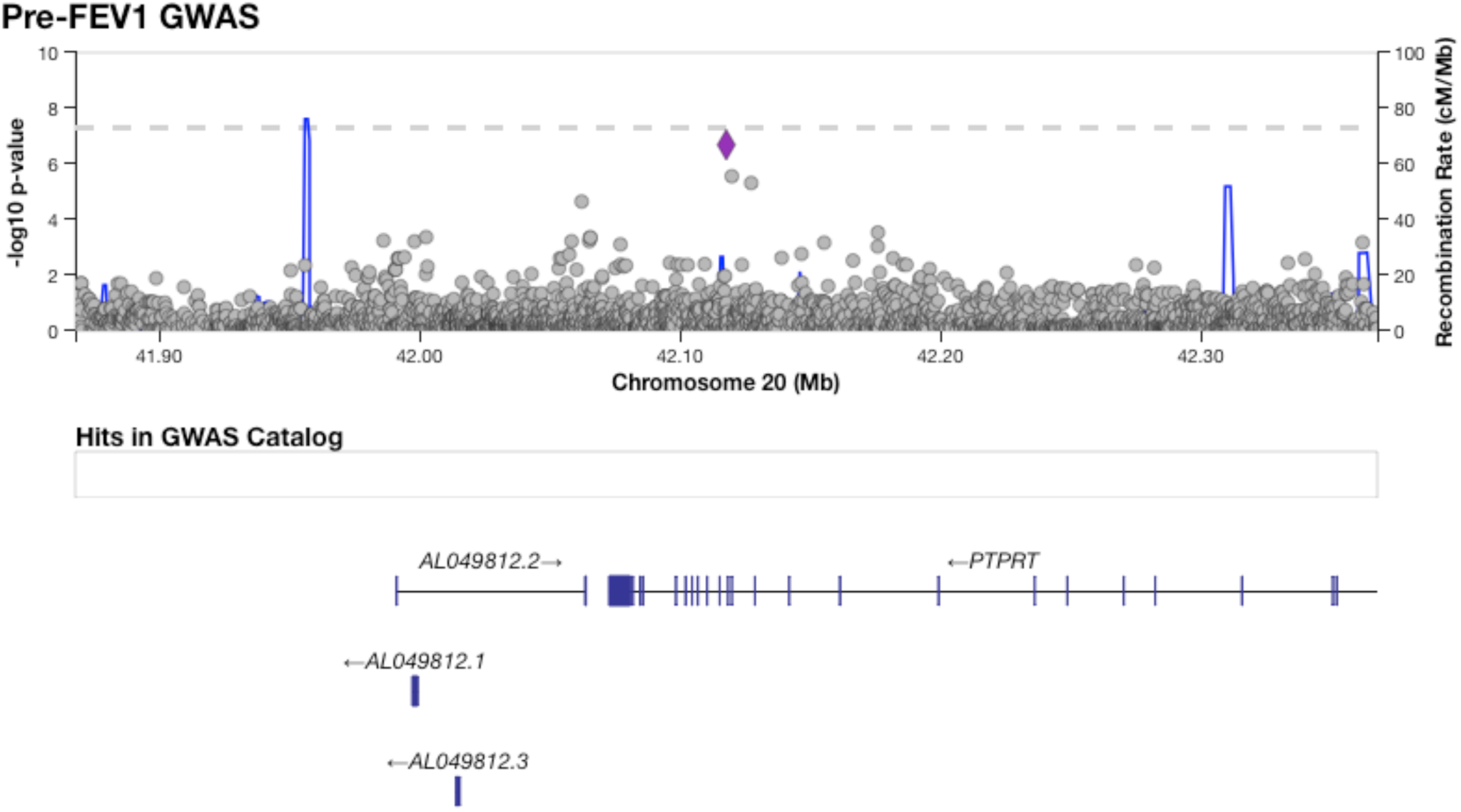
The region around a variant on chromosome 20 suggestively associated with Pre-FEV_1_.

**Supplementary Figure 15:**
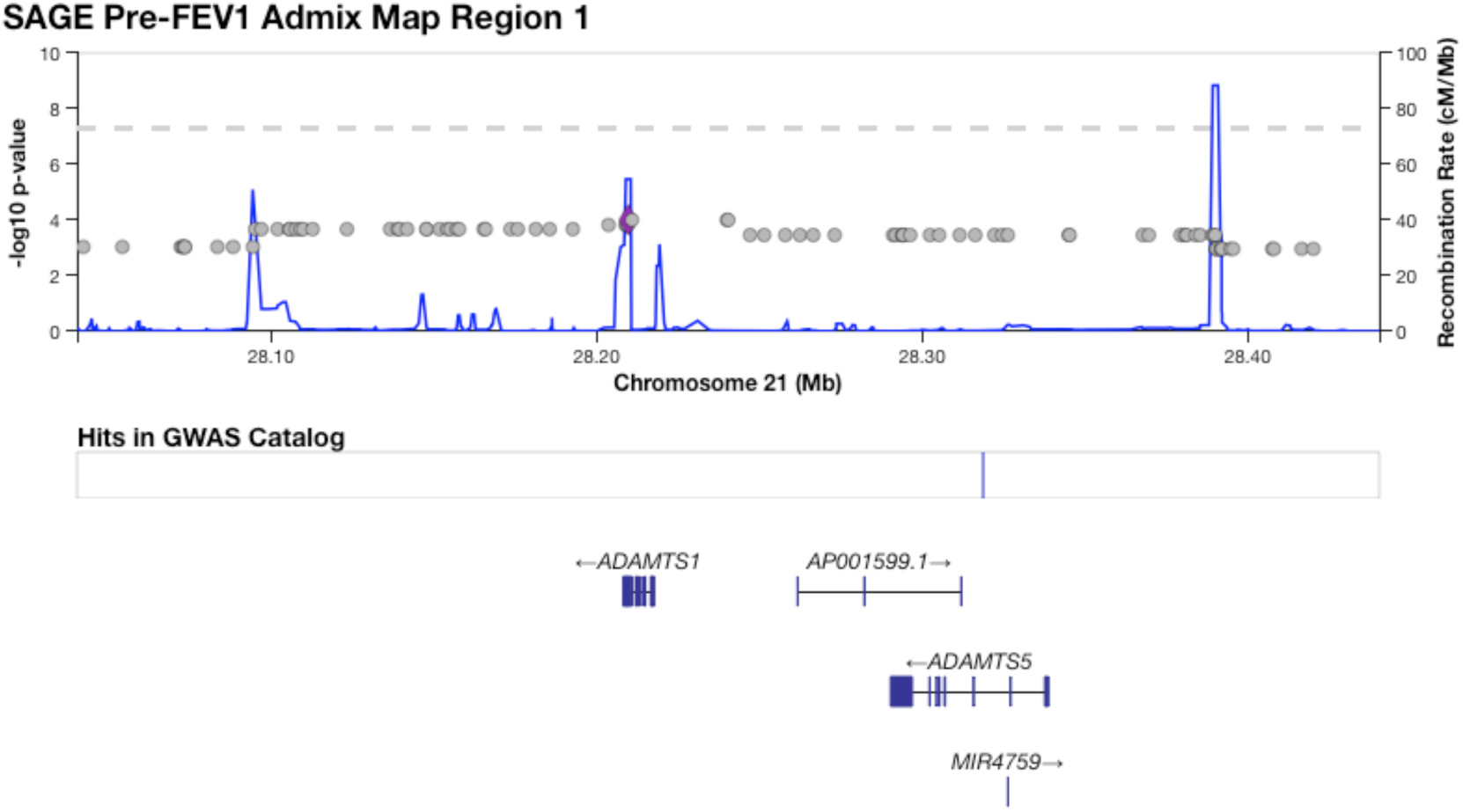
The region on chromosome 21 identified by admixture mapping as significantly associated with Pre-FEV_1_.

**Supplementary Figure 16:**
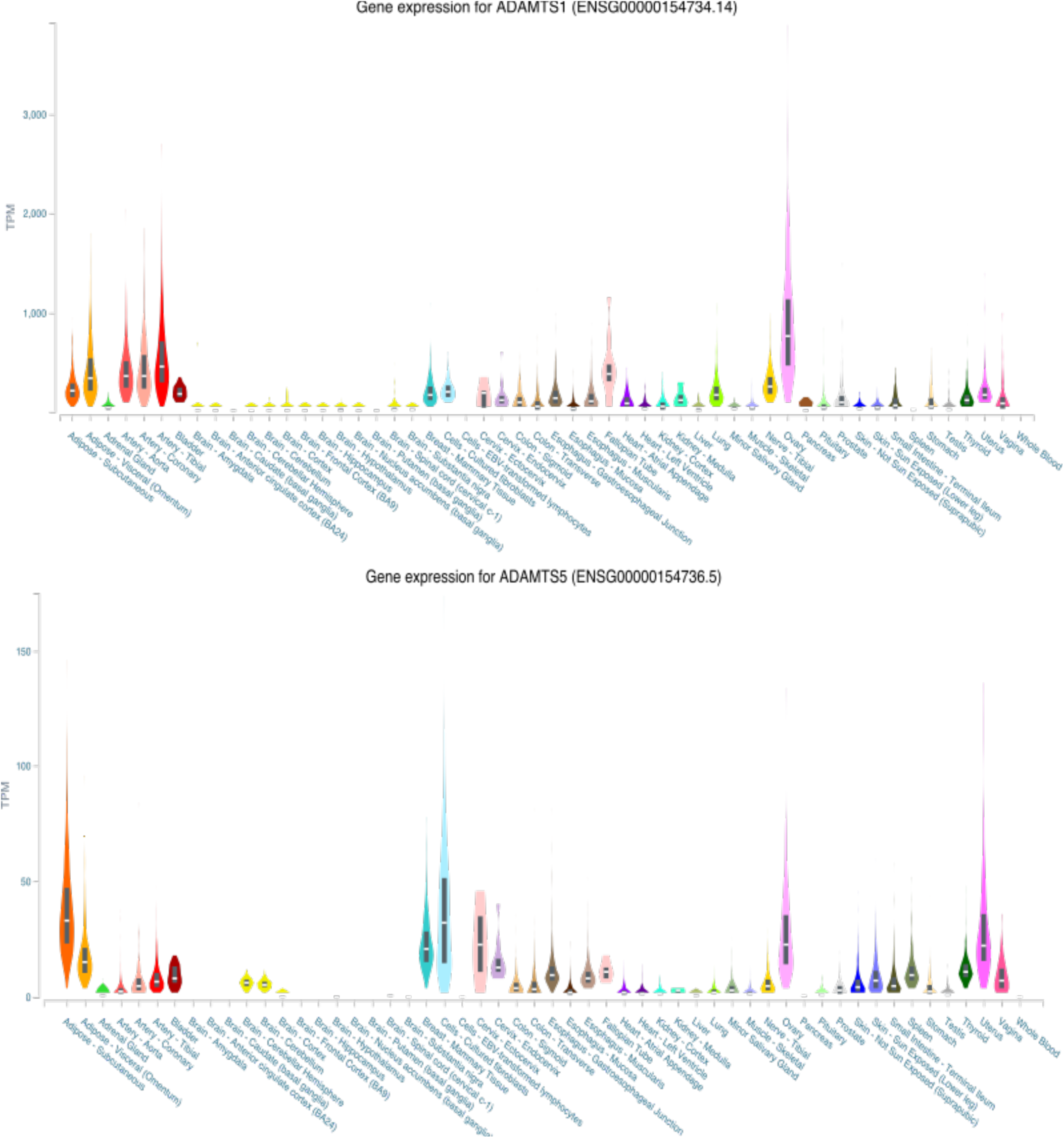
Tissue-specific gene expression for *ADAMTS1* (top) and *ADAMTS5* (bottom) from GTEx v8. Both genes are expressed in certain tissues, such as ovary and vagina. However, *ADAMTS1* shows markedly higher expression in lung and arterial tissue, while *ADAMTS5* is more strongly expressed in breast, fibroblast, cervix, adipose, and whole blood.

**Supplementary Figure 17:**
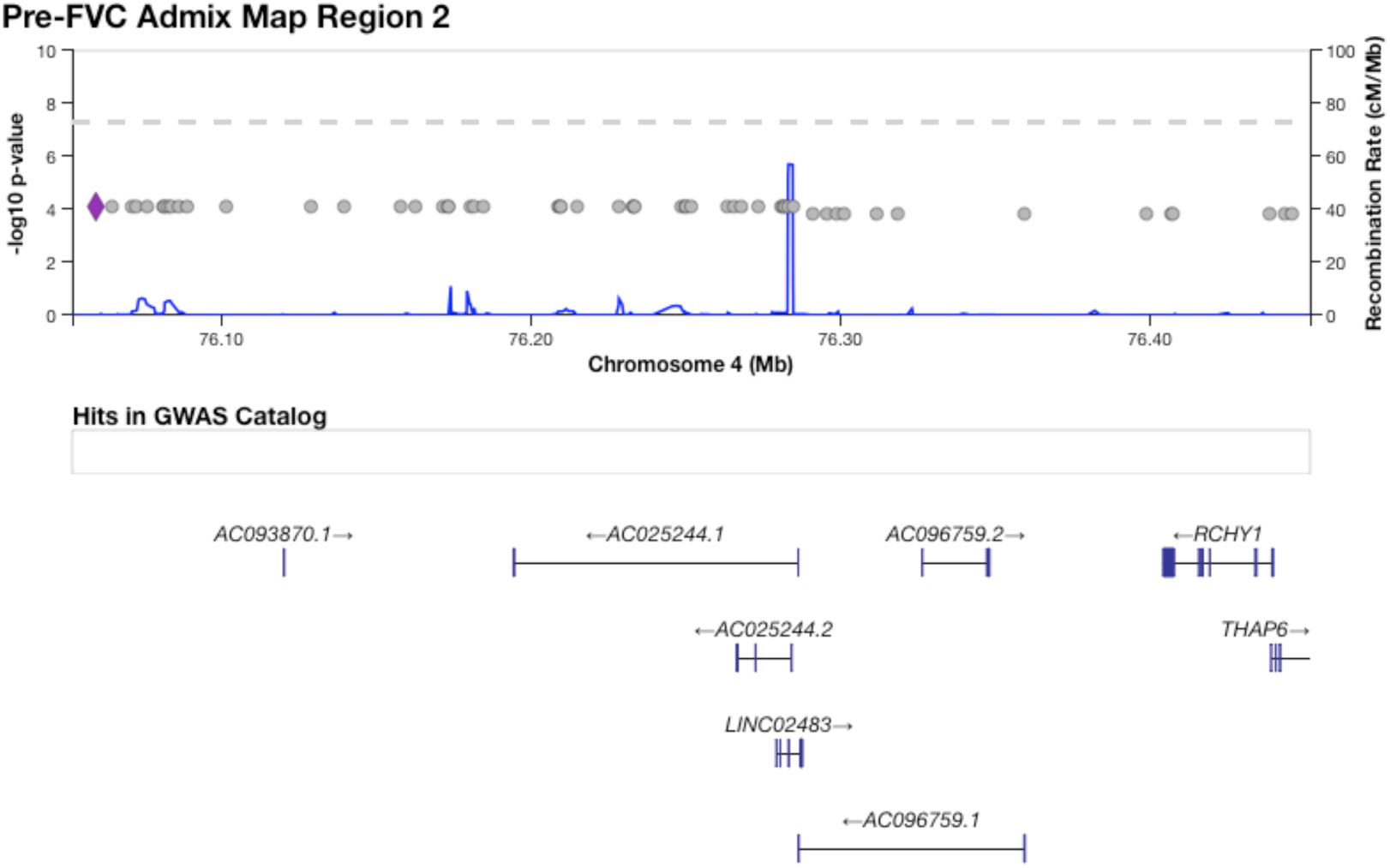
The region on chromosome 4 identified by admixture mapping as significantly associated with Pre-FVC.

**Supplementary Figure 18:**
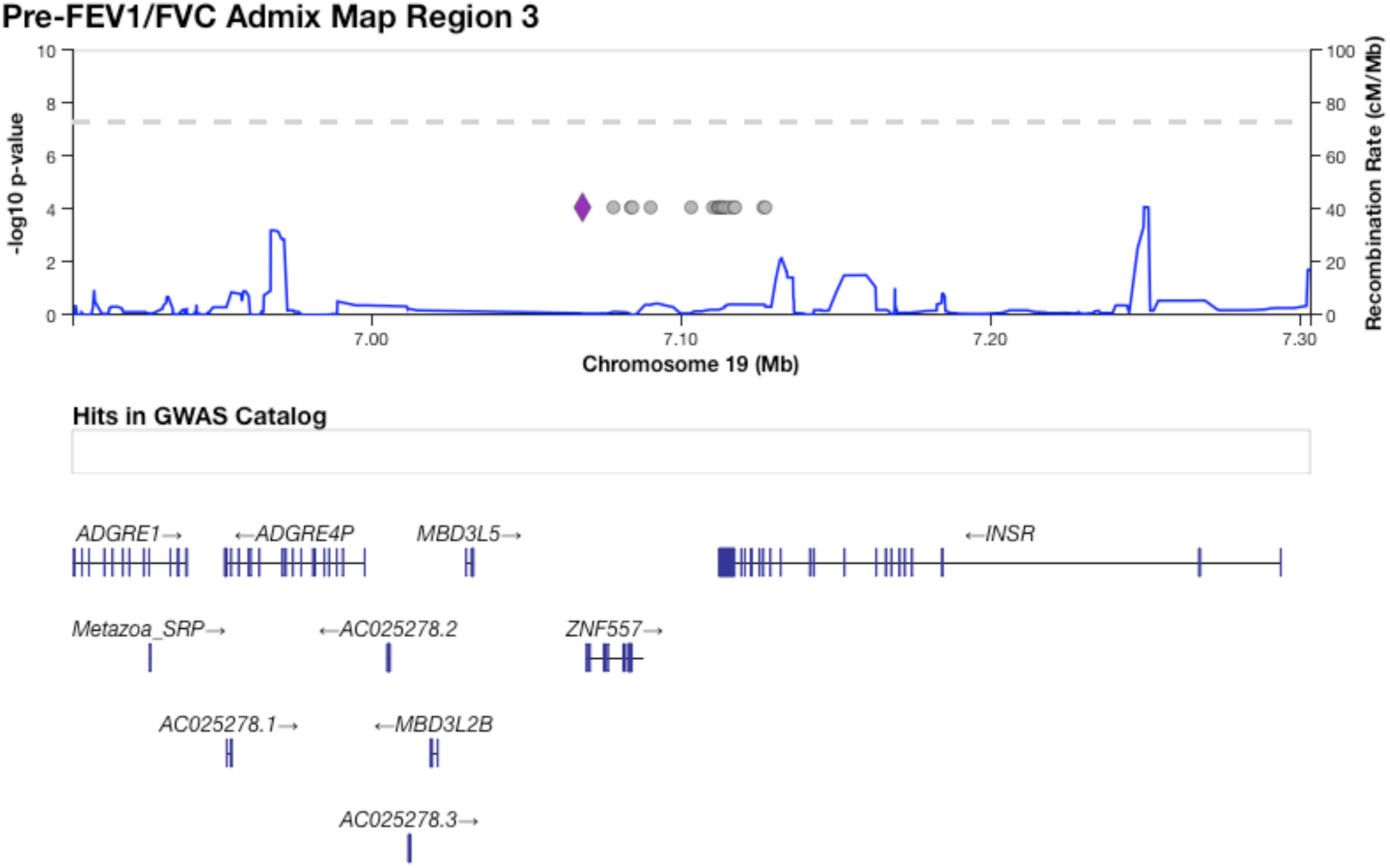
The region on chromosome 19 identified by admixture mapping as significantly associated with Pre-FEV_1_/FVC.

**Supplementary Figure 19:**
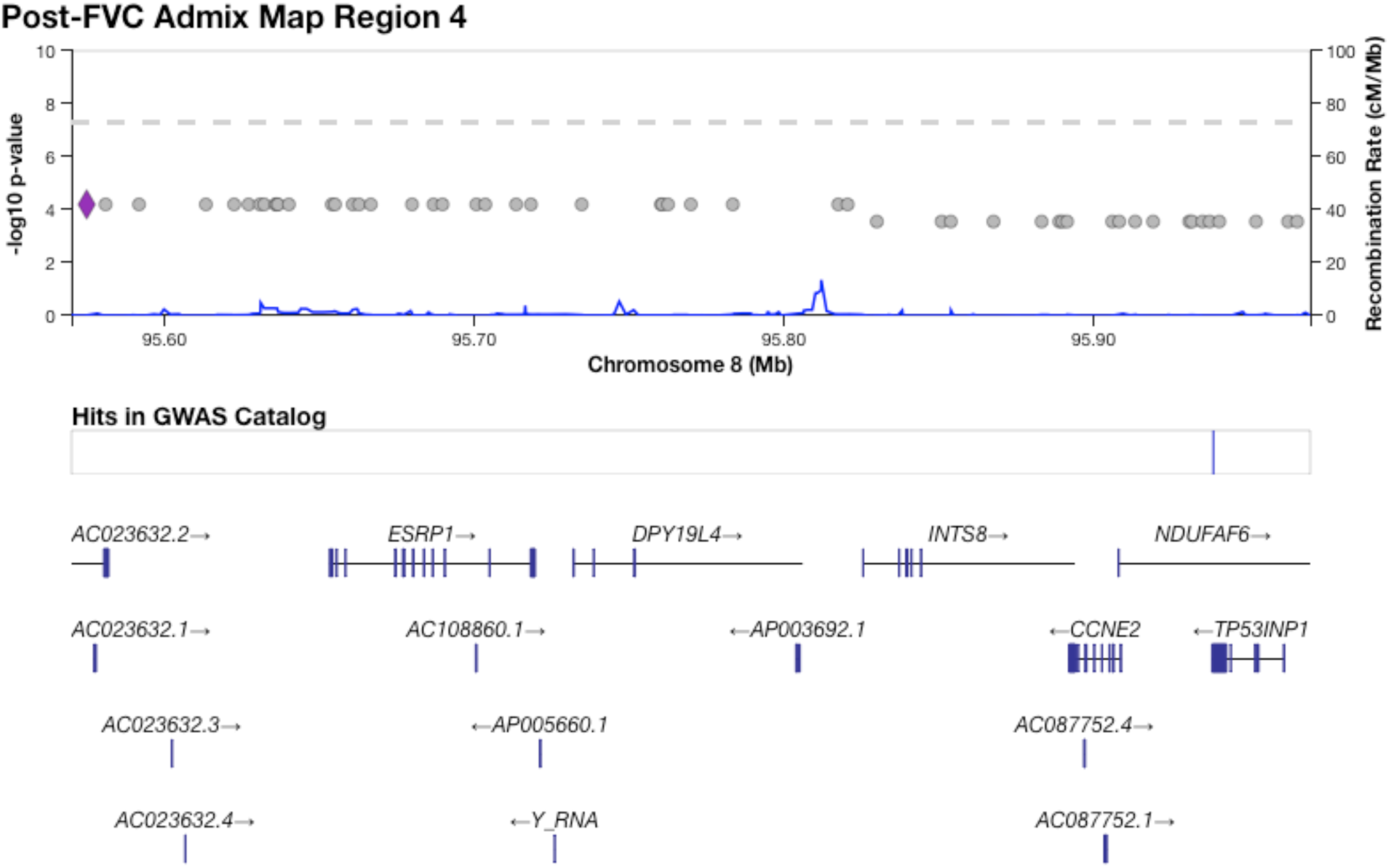
The region on chromosome 8 identified by admixture mapping as significantly associated with Post-FVC.

**Supplementary Figure 20:**
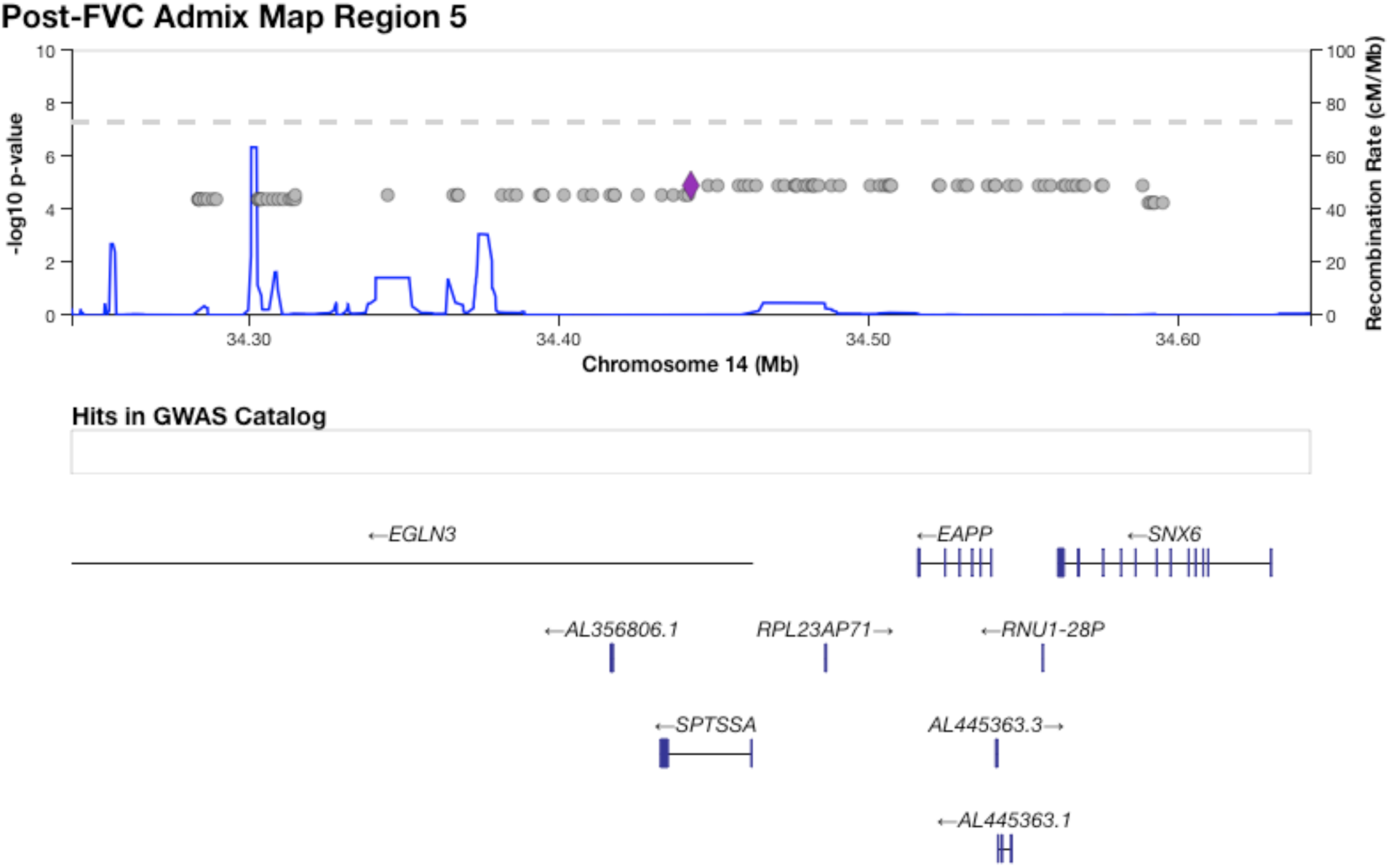
The region on chromosome 14 identified by admixture mapping as significantly associated with Post-FVC.

**Supplementary Figure 21:**
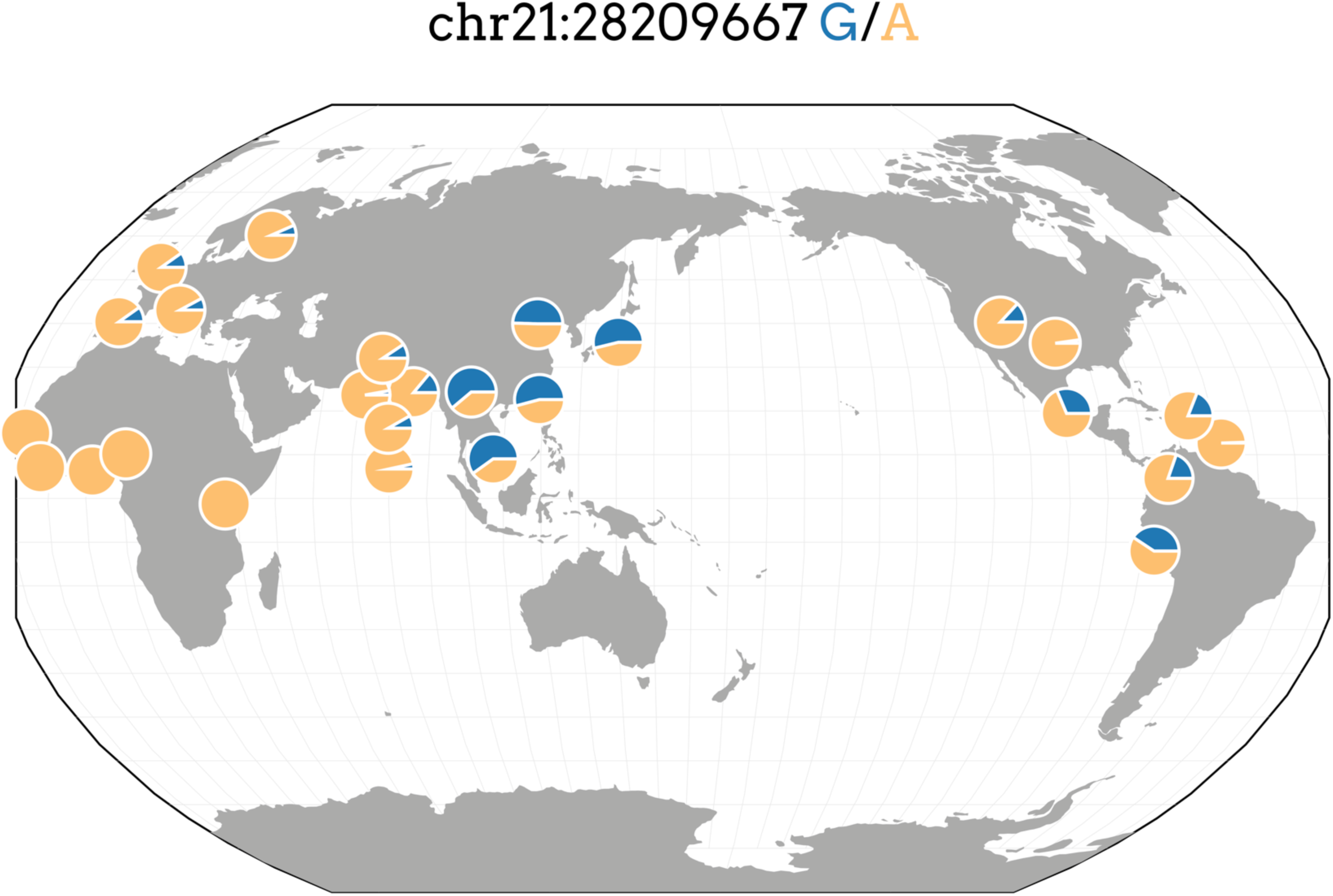
Distribution of minor allele frequencies for SNP rs13615 in 1000 Genomes populations (hg19 build). The derived allele G shows strong separation by distance, with high prevalence in East Asia, moderately high prevalence in the Americas, and low prevalence in Europeans and South Asians. However, in Africans and African-derived populations from 1000 Genomes, the G allele occurs at very low frequency, if at all.

**Supplementary Figure 22:**
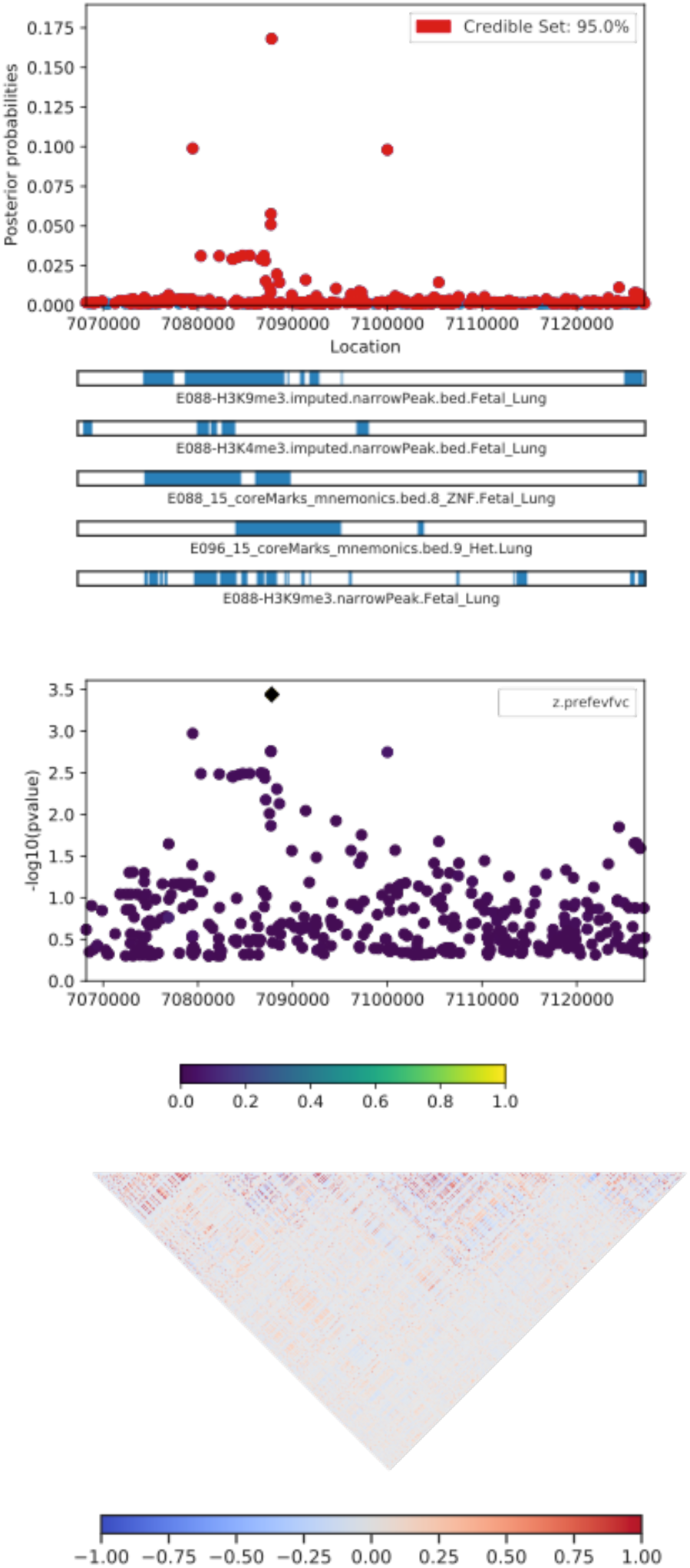
A CANVIS plot of results from PAINTOR fine-mapping for locus 3, an association on chromosome 19 with Pre-FEV_1_/FVC. The SNP with highest posterior probability of causality (0.168) is rs72986681.

**Supplementary Figure 23:**
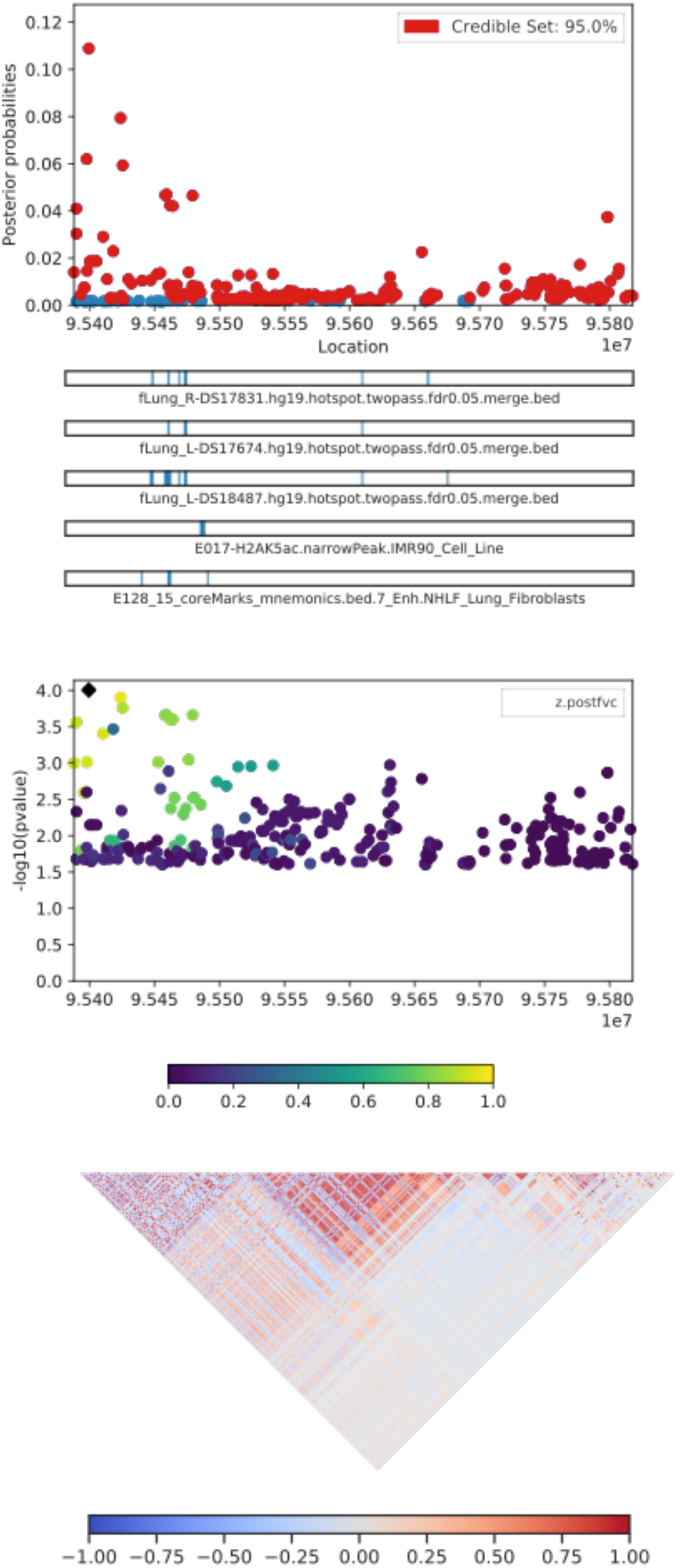
A CANVIS plot of results from PAINTOR fine-mapping for locus 4, an association on chromosome 8 with Post-FVC. The SNP with highest posterior probability of causality (0.109) is rs2470740.

**Supplementary Figure 24:**
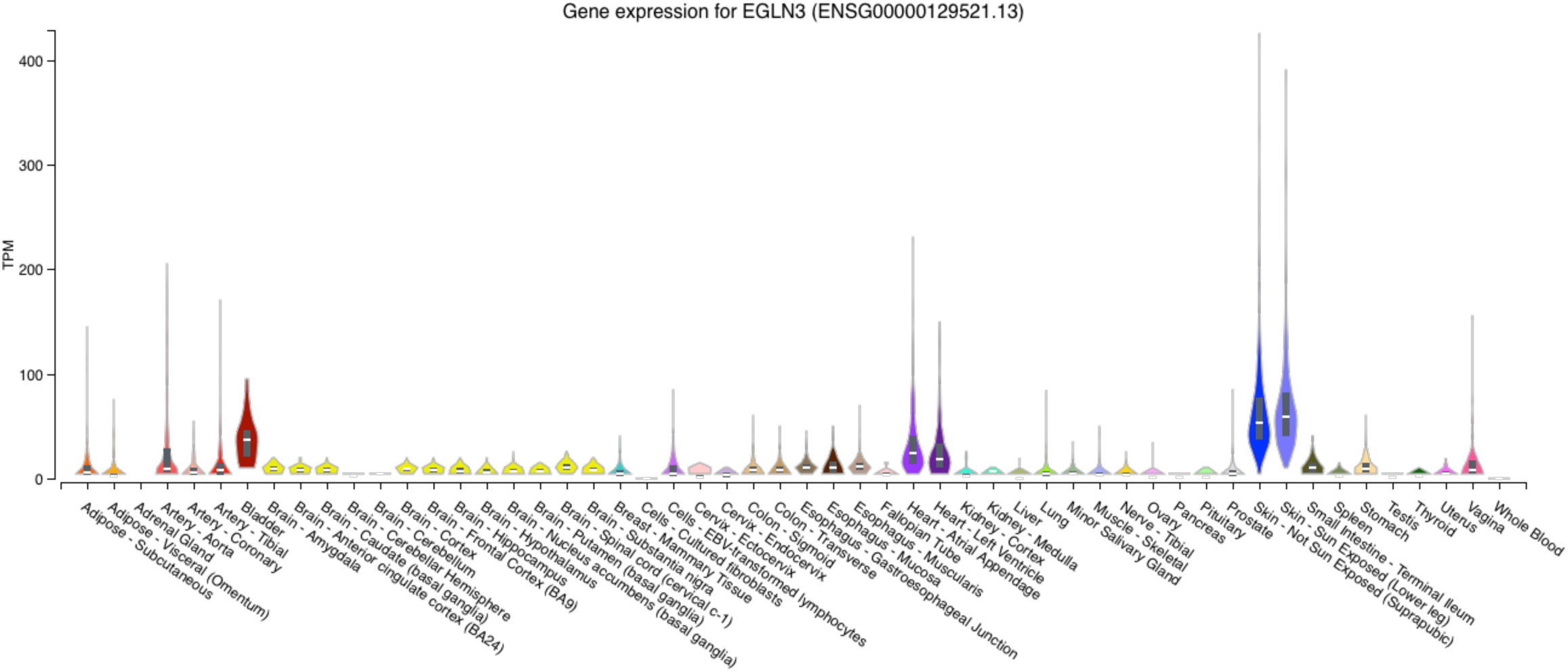
Expression profile for the gene *EGLN3* across tissue types from GTEx v8. EGNL3 is most strongly expressed in skin and bladder, but also shows nontrivial expression in heart and arterial tissue.

**Supplementary Figure 25:**
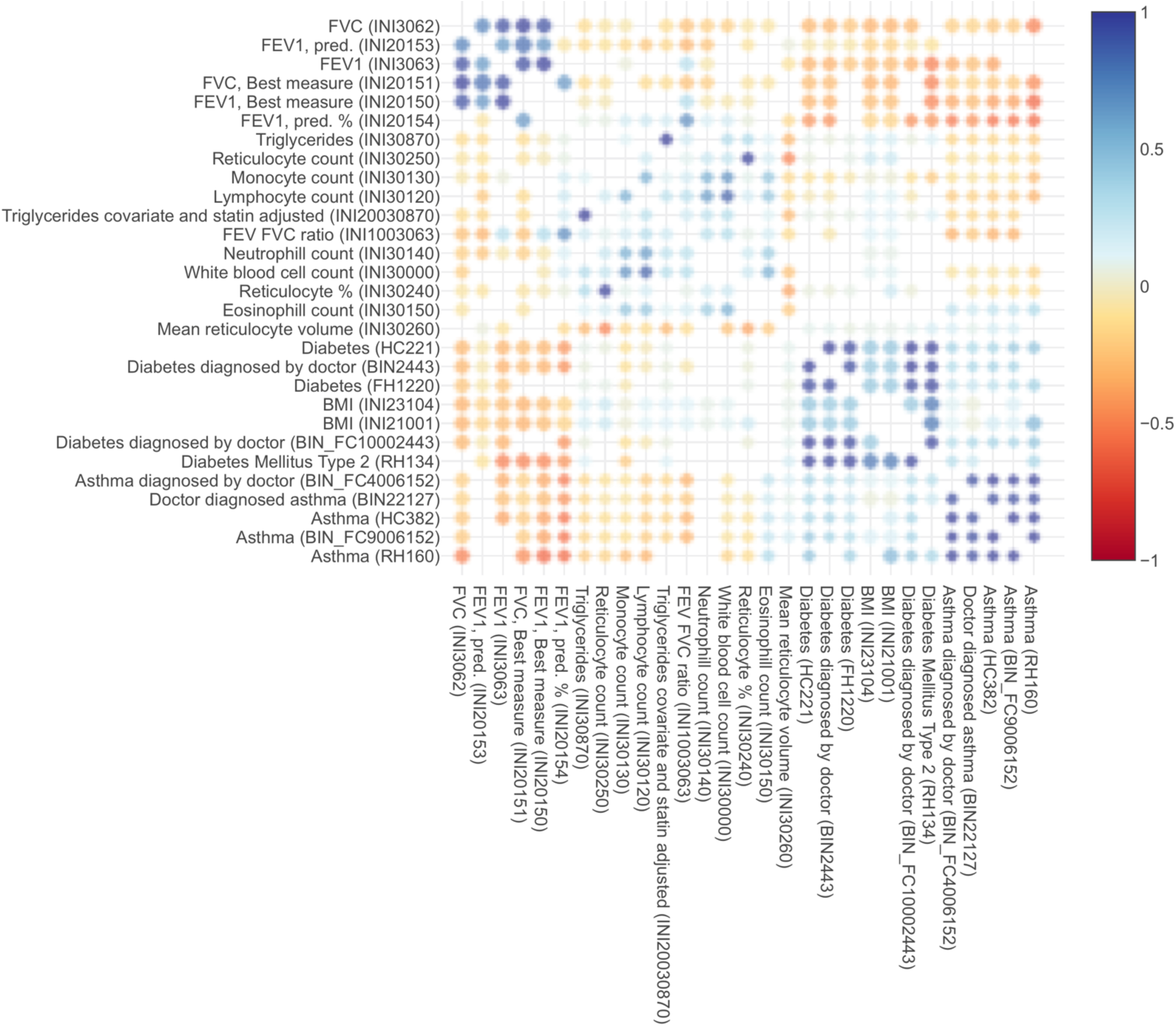
Genetic correlations between lung, heart, blood, and obesity traits, computed from the UK Biobank.

